# Complex introgression among three diverged largemouth bass lineages

**DOI:** 10.1101/2021.05.12.443886

**Authors:** Katherine Silliman, Honggang Zhao, Megan Justice, Wilawan Thongda, Bryant Bowen, Eric Peatman

**Affiliations:** School of Fisheries, Aquaculture, and Aquatic Sciences, Auburn University, Auburn, AL 36849, USA; Department of Natural Resources, Cornell University, Ithaca, NY, 14850, USA; Center of Excellence for Shrimp Molecular Biology and Biology (CENTEX Shrimp), Faculty of Science, Mahidol University, Rama VI Road, Bangkok, 10400, Thailand; National Center for Genetic Engineering and Biotechnology (BIOTEC), National Science and Technology Development Agency (NSTDA), Pathum Thani, 12120, Thailand; Georgia Department of Natural Resources, Social Circle, GA, 30025, USA

**Author notes:** Address for correspondence: Eric Peatman, School of Fisheries, Aquaculture, and Aquatic Sciences, Auburn University, Auburn, AL 36849, USA.

**Keywords:** largemouth bass, hybridization, Mobile-Tensaw Delta, single nucleotide polymorphisms, genotype-by-sequencing

## Abstract

Hybrid zones between diverged lineages offer an unique opportunity to study evolutionary processes related to speciation. Natural and anthropogenic hybridization in the black basses (*Micropterus* spp.) is well documented, including an extensive intergrade zone between the widespread northern Largemouth Bass (*M. salmoides*) and the Florida Bass (*M. floridanus*). Phenotypic surveys have identified an estuarine population of Largemouth Bass (*M. salmoides*) in the Mobile-Tensaw Delta, with larger relative weight and smaller adult size compared to inland populations, suggesting a potential third lineage of largemouth bass. To determine the evolutionary relationships between these Mobile Delta bass populations, *M. salmoides*, and *M. floridanus*, putative pure and intergrade populations of all three groups were sampled across the eastern United States. Phylogenetic analyses of 8,582 nuclear SNPs derived from genotype-by-sequencing and the ND2 mitochondrial gene determined that Delta bass populations stem from a recently diverged lineage of Largemouth Bass. Using a novel quantitative pipeline, a panel of 73 diagnostic SNPs was developed for the three lineages, evaluated for accuracy, and then used to screen 881 samples from 52 sites for genetic integrity and hybridization on the Agena MassARRAY platform. These results strongly support a redrawing of native ranges for both the intergrade zone and *M. floridanus*, which has significant implications for current fisheries management. Furthermore, Delta bass ancestry was shown to contribute significantly to the previously described intergrade zone between northern Largemouth Bass and Florida Bass, suggesting a more complex pattern of secondary contact and introgression among these diverged *Micropterus* lineages.

## Introduction

Hybrid zones are geographical areas of ongoing hybridization between diverged lineages, and therefore offer a unique opportunity to study evolutionary processes related to speciation (Barton & Hewitt, 1985; Gompert et al., 2017). The most accepted process by which hybrid zones arise is secondary contact of diverged lineages that have not evolved complete reproductive isolation. This contact may occur due to natural shifts in geological or environmental barriers, or anthropogenic introductions of non-native species (Largiadèr, 2007; Woodruff, 1973). Nearly all studied hybrid zones are those involving two parental taxa, however more complex hybrid zones involving multiple taxa are known to occur in nature (Chhatre et al., 2018; Keck & Near, 2009; Nevado et al., 2011; Peñaloza-Ramírez et al., 2010). When hybridization is thought to be the result of anthropogenic translocations of nonnative species, it can be of particular concern due to loss of biodiversity (McDonald et al., 2008; Rhymer & Simberloff, 1996; Viard et al., 2020).

Interspecific hybridization is relatively common among freshwater fishes. Of over 150 described pairs of hybridizing species in the US, 20% are found within Centrarchidae, including the black basses (*Micropterus* spp.) (Bolnick, 2009). Natural and anthropogenic hybridization in black basses is well documented, using either morphometrics (Bailey & Hubbs, 1949; Baker et al., 2013) or diagnostic genetic markers (Bagley et al., 2011; Bangs et al., 2018; Barthel et al., 2010; C. Li et al., 2015; Lutz-Carrillo et al., 2006; Philipp et al., 1983; Thongda et al., 2020). One of the most notable cases of hybridization in *Micropterus* is an extensive intergrade zone between the widespread northern Largemouth Bass (*M. salmoides*) and the Florida Bass (*M. floridanus*) (see Note), of which pure populations are thought to be restricted to peninsular Florida. This hybrid zone was initially described using morphology and meristics, and was thought to extend through northern Florida, Georgia, and small parts of Alabama and South Carolina (Bailey & Hubbs, 1949). Thirty years later, (Philipp et al., 1983) evaluated allele frequencies at two diagnostic allozymes which extended the hybrid zone west to Mississippi and north to Virginia. Their result is unsurprising, for disconnects between morphological and genetic descriptions of hybrid zones are common, as phenotype can often be inaccurate for identifying later-generation hybrids (Meyer et al., 2017; A. T. Taylor et al., 2018). While Philipp et al. (1983) acknowledged the inaccuracy of meristics for detecting hybrids, they also suggested that the apparent hybrid zone expansion was due to stockings of Florida Bass outside their native range for recreational fishing. This latter view has persisted, with a recent review portraying the geographic extent of natural hybridization the same as was first described by Bailey and Hubbs (A. T. Taylor et al., 2019). Modern genetic tools are required to accurately characterize the hybrid zone of *M. salmoides–M. foridanus* and determine whether this zone is primarily shaped by stocking or historical hybridization.

In addition to the high diversity of *Micropterus* species, the southeastern United States has the greatest aquatic diversity in North America (Warren et al., 1997), which is exemplified by the numerous freshwater and brackish-water fish species supported by the Mobile-Tensaw River Delta (Swift et al., 1986; H. A. Swingle & Bland, 1974; W. E. Swingle et al., 1966). Due to a rich history of geologic shifts, glaciation, and sea level fluctuations, freshwater fishes of the southeastern U.S. have experienced repeated bouts of habitat isolation and reconnection (Swift et al., 1986). These dramatic changes in recent evolutionary history are likely responsible for the region’s high endemicity and hybridization observed in numerous freshwater fish, such as mosquitofishes (Wilk & Horth, 2016), crappies (Travnichek et al., 1996), sunfish (Avise & Saunders, 1984), and bass (Near et al., 2003). Such sea level fluctuations in the Late Pliocene are thought to have isolated *M. floridanus* from *M. salmoides* on the Florida peninsula (Near & Kim, 2021), resulting in genetically and phenotypically diverged sister taxa.

In the Mobile Delta exists another phenotypically distinct population of largemouth bass. These “Delta” bass have a smaller adult size and higher relative weight compared to other *M. salmoides* populations, as well as physiological differences that may facilitate their survival in water with elevated salinity (up to 13 ppt) (DeVries et al., 2015; Glover et al., 2012, 2013; Tucker, 1985). Preliminary genetic analysis using isozymes and microsatellites showed that Delta bass were genetically similar to *M. salmoides*, but were unable to conclusively determine if they were a distinct genetic lineage (DeVries et al., 2015; Hallerman et al., 1986). Given the proximity of the Mobile Delta to the *M. salmoides–M. floridanus* intergrade zone, a thorough genetic characterization of Delta bass is necessary for determining how this putative lineage contributes to existing hybridization patterns.

The objectives of this study were to employ modern genomic techniques to analyze the *M. salmoides–M. floridanus* hybrid zone and resolve the phylogenetic relationship of Largemouth Bass from the Mobile-Tensaw Delta. Specifically, we 1) used thousands of single nucleotide polymorphisms (SNPs) and mitochondrial sequencing to characterize the phylogenetic and population genetic relationships between Delta Largemouth Bass, northern Largemouth Bass, and Florida Bass, 2) developed a SNP assay for accurate and rapid identification of pure and hybrid individuals, 3) applied this SNP assay to hundreds of samples across the eastern U.S. in order to characterize the geographic extent of hybridization and assess the role of stocking in driving the extent of Florida Bass introgression.

## Methods

### Sample collection and GBS library preparation

Sample collection and genotype-by-sequencing (GBS) using the *Pst*I restriction enzyme was previously described in Thongda et al. (2019). For the present study, we used GBS data for 144 largemouth bass samples across 18 sites and captive hatchery populations: Largemouth Bass from the Mobile-Tensaw Delta (DLB) (*M. salmoides*, N = 29), northern Largemouth Bass (NLB) (*M. salmoides*; N = 42), Florida Bass (FLB) (*M. floridanus*; N = 29), and putative intergrade largemouth bass (ILB) (N = 44) (Supp. Table 1). For phylogenetic analysis, we also included 17 outgroup black bass samples: Spotted Bass (*M. punctulatus*; n = 2), Coosa Redeye Bass (*M. coosae*; 5), Shoal Bass (*M. cataractae*; n = 5), Smallmouth Bass (*M. dolomieu*; 3), and Guadalupe Bass (*M. treculii*; n = 2). FLB, NLB, and ILB individuals were identified using previously developed 25-38 SNP panels that are diagnostic for northern Largemouth and Florida Bass (C. Li et al., 2015; Zhao et al., 2018). We also sampled an additional 786 largemouth bass individuals across 52 populations for extensive population genetic structure analysis and hybrid classification using the diagnostic SNP panel developed in this study (Fig. 1; also refer to online Supplementary material, File 2). For the 881 total samples genotyped with the diagnostic SNP panel, DNA was extracted from fin clips using a simple sodium hydroxide (NaOH) and hydrochloric acid (HCl) tissue digestion (Truett et al., 2000). For mitochondrial gene sequencing, DNA was extracted from fin clips or whole blood samples using E.Z.N.A. DNA Kits (Omega BioTek, Melbourne, Australia).

**Figure 1.**
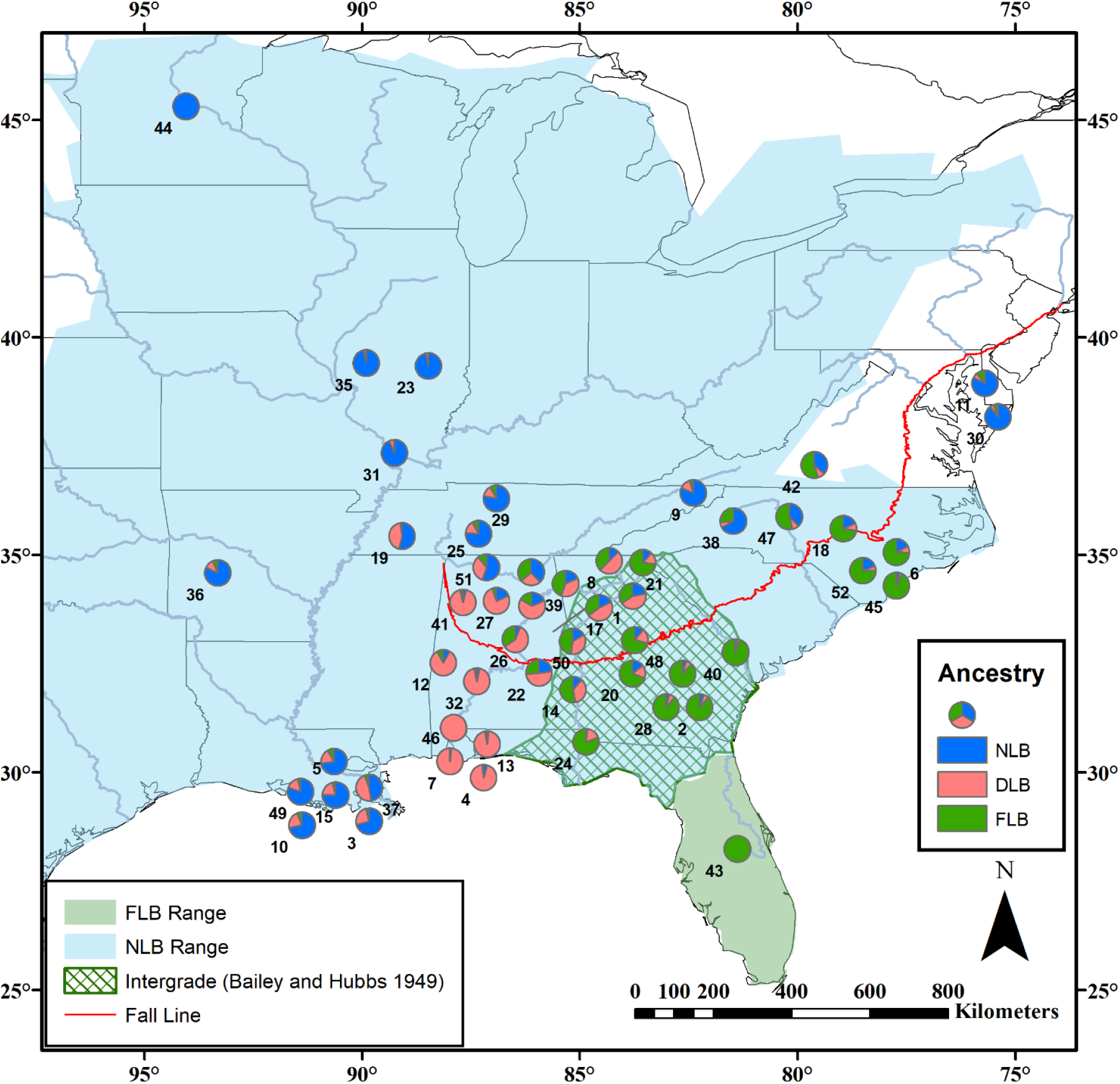
Sampling sites of largemouth bass assayed with 73 SNP panel. Populations are labelled as in Table S2 and represented by pie graphs showing the mean estimated ancestry proportions for three bass lineages based on 73 diagnostic SNPs and the population genetic program STRUCTURE (K=3). Delta Largemouth Bass (DLB) are pink, northern Largemouth Bass (NLB) are blue, and Florida Bass (FLB) are green. Native ranges of NLB and FLB are shown, as well as the NLB–FLB hybrid zone as described by Bailey & Hubbs (1949). Map data from ESRI; FLB and NLB native ranges from Taylor et al. (2019).

### GBS analysis and SNP calling

Raw GBS reads were demultiplexed, quality filtered, and aligned to a *M. floridanus* genome reference (N50 of 11,136 bp and 249,768 scaffolds (Zhao et al., 2018)) using STACKS v2.55 (Rochette et al., 2019). GBS reads were trimmed and de-multiplexed using the process_radtags program in STACKS, while removing reads with an uncalled base (-c) and low-quality scores (-q). For reference-based SNP calling, we mapped the de-multiplexed reads to the *M. floridanus* genome using the Burrows–Wheeler Aligner v0.7.17 with default settings (H. Li & Durbin, 2009). The mapped reads were sorted using the “sort” function in SAMTOOLS v1.6 (Heng Li et al., 2009), then used to call SNPs with the “gstacks” pipeline (-var-alpha 0.05, -gt-alpha 0.05, --max-clipped 0.3). The “populations” program was used to merge loci that were produced from the same restriction enzyme cut sites (merge sites), perform initial SNP filtering, and export a variant call format (VCF) file. Additional SNP filtering was done with VCFtools v0.1.17 (Danecek et al., 2011), custom Python code, and code adapted from Jon Puritz’s laboratory (Puritz et al., 2014).

For population genetic analyses, biallelic SNPs were retained if they had a minor allele frequency greater than 0.05, had a minor allele count greater than 4, were genotyped in at least three population groups (NLB, DLB, FLB, or ILB), and had less than 15% missing data across all individuals. SNPs that departed Hardy-Weinberg equilibrium with a p-value cutoff of 0.05 in two or more sampling sites were also removed. SNPs in linkage disequilibrium (r^2^ > 0.2) or within the same 1000 bp window were pruned using BCFtools +prune plugin (Heng Li et al., 2009). For phylogenetic analyses that included outgroup *Micropterus* species, variable sites were retained if they were genotyped in at least 4 species and 70% of all individuals, were heterozygous in fewer than 70% of samples (--max-obs-het), and had a minor allele count greater than 3. The VCF output of stacks was converted to a phylip file using vcf2phylip v2.0 (Ortiz, 2019).

### ND2 sequencing

DNA from 62 samples were amplified with previously published primers targeting the mitochondrial NADH subunit 2 (ND2) gene (Kocher et al., 1995). Amplification of the ND2 gene used 25 ml reaction volumes and included 1X Kapa HiFi HotStart ReadyMix (2X) (Kapa Biosystems, Wilmington, MA), 0.2 mM forward primer, 0.2 mM reverse primer, and 2 μL of template DNA. All PCRs included negative control reactions (no DNA template). Thermal cycler reactions were conducted with a heated lid and the following conditions: initial denaturation at 95°C for 3 min, followed by 5 cycles at 94°C for 30 s, annealing at 57°C for 30 s, and 72°C for 75 s; an additional 5 cycles of annealing at 56°C for 30 s and 5 cycles with annealing at 55°C for 30 s; followed by 20 cycles at 94°C for 30 s, annealing at 54°C for 30 s and 72°C for 75 s, and a final extension at 72°C for 10 min., followed by a final hold at 4°C (Baker et al., 2013). PCR products were visualized on a 1% agarose gel to check the amplification success and that a single band was observed. Amplicons were then cleaned using ExoSAP-IT (ThermoFisher Scientific, Waltham, MA). Quantity and quality of PCR products was determined with a NanoDrop Spectrophotometer (ThermoFisher Scientific, Waltham, MA).

Sanger cycle sequencing of all products was performed by Genewiz, Inc. (North Plainfield, NJ) using the PCR primers and internal ND2 primer MET-F (Kocher et al., 1995). Consensus sequences of ND2 were obtained by analyzing the forward sequencing trace files using CodonCode Aligner (2019.2.1), then trimmed to the first 598 bp. The reads were inspected manually for quality and for any discrepancies in base calls. All sequences were aligned in MEGA-X using MUSCLE to determine the number of unique haplotypes present for subsequent analyses (Edgar, 2004; Kumar et al., 2018). A minimum spanning network for haplotypes was calculated and visualized in the R package pegas v0.14 (Paradis, 2010).

### Phylogenetics and population genetics

Maximum likelihood phylogenetic analyses of 100 individuals from pure NLB, FLB, and DLB populations and 17 outgroup *Micropterus* samples were conducted using 18,492 concatenated variable GBS sites with IQ-TREE v1.6.12 (Nguyen et al., 2015). ModelFinder as implemented in IQ-TREE used BIC and determined the best-fit substitution model that included ascertainment bias correction to be TVM+F+ASC+R3 (Kalyaanamoorthy et al., 2017). Branch supports were assessed with both 1000 ultrafast bootstrap approximation replicates and 1000 bootstrap replicates for the SH-like approximate likelihood ratio test (Guindon et al., 2010; Hoang et al., 2018). The resulting tree was rooted with the five outgroup bass species. The same analysis was performed on 18,641 SNPs with ILB individuals included. The resulting best trees were plotted with the R package ggtree (Yu et al., 2017).

Population genetic summary statistics were calculated on GBS SNPs for population groups (NLB, DLB, FLB, ILB) and individual sampling sites with at least 5 individuals. Observed heterozygosity (H_o_), expected heterozygosity (H_e_), overall F_ST_, and F_IS_ were calculated using the basic.stats function in the R package hierfstat v0.5.7 (Goudet & Jombart, 2015). Confidence intervals for population-specific F_IS_ were determined using the boot.ppfis function in hierfstat with 1,000 bootstrap replicates. Pairwise F_ST_ following Weir and Cockerham (1984) was calculated using the genet.dist function in hierfstat (Weir & Cockerham, 1984). Variable sites were determined to be “fixed” between groups if one group had an allele frequency ≥ 0.98 in one group and < 0.02 in the other. This cutoff was chosen to allow for singleton alleles observed in a lineage that may be due to genotyping error.

The model-based Bayesian clustering method STRUCTURE v2.2.4 (Pickrell & Pritchard, 2012) was used to determine the number of distinct genetic clusters (K) with a burn-in period of 50,000 repetitions followed by 200,000 repetitions. Five replicate analyses were performed on the thinned GBS SNP dataset with values of K = 1–10. Replicates were summarized and visualized using the CLUMPAK server (Kopelman et al., 2015). The ΔK method implemented in STRUCTURE HARVESTER was used to determine an optimal K (Earl & vonHoldt, 2012). Principal component analysis (PCA) was implemented in the R package adegenet v2.1.1 (Jombart & Ahmed, 2011) and visualized with the R package PCAviz v0.3.29 (Novembre et al., 2018). Missing data were filled by randomly drawing an allele based on the overall allele frequency across all individuals in a group using custom R code.

### Panel Development, Testing, and Genotyping

Our goal when developing a SNP assay for accurate identification of largemouth bass lineages and hybrids was to find a subset of SNPs that best matched the ancestry results determined by STRUCTURE when using the full GBS SNP dataset. We first used custom Python scripts and the populations.sumstats.tsv output from Stacks to identify 2,809 SNPs that were diagnostic between DLB, NLB, and FLB, with the criteria that they had > 90% or < 10% allele frequency in at least two of the three largemouth bass lineages. PCA was then performed on these diagnostic SNPs for 100 ‘pure’ largemouth bass samples (excluding intergrades) as previously described. To identify SNPs that had the greatest contributions to the PCA and therefore were the most diagnostic, we extracted 549 SNPs in the 90% quantile of loadings for PC1 and PC2. We then randomly sampled 80-100 SNPs from this set for analysis with STRUCTURE and compared the Q-values for each individual with the Q-values derived using the full SNP dataset. This analysis was iteratively repeated 300 times. A subset of 127 total SNPs with the fewest samples exhibiting a > 5% difference in Q-values was retained for marker development.

A MassARRAY System (Agena Bioscience, San Diego, California) was used to develop, genotype, and evaluate SNP panels. For each of the 127 diagnostic SNPs, a 201 bp sequence was extracted from the *M. floridanus* genome (the SNP and 100 bp flanking on either side) and imputed into the MassARRAY Assay Design Software to design two multiplex assays with a maximum of 60 SNPs per assay. This produced forward, reverse, and extension primer sequences for assays of 52 and 39 SNPs (Supp. File 3), which were ordered through IDT (Integrated DNA Technologies, Inc.). We used 91 samples to test the concordance of SNP genotypes generated with the MassARRAY platform and GBS, as well as 70 technical replicates made up of 40 individuals to assess the consistency of genotype calls between runs. Discordant genotypes due to missing data were excluded from this analysis. SNPs that consistently failed in greater than 90% of samples, were invariant, or inconsistent between runs were removed, resulting in a final set of 73 SNPs. For all samples run on the MassARRAY platform, amplification and extension reactions were performed using the iPLEX Gold Reagent Kit (Agena Bioscience, San Diego, California) according to the manufacturer’s protocol (Gabriel et al., 2009). SNP genotypes were called using the MassARRAY Typer 4 analysis software and manually confirmed. This software uses a three-parameter (mass, peak height, and signal-to-noise ratio) model to estimate genotype probabilities. Genotype concordance and SNP panel evaluation was performed with custom Python scripts and Excel.

Some samples used for GBS, as well as 786 additional samples across 52 sites, were genotyped at these 73 SNPs for a total of 881 samples. To determine the FLB, NLB, and DLB ancestry proportions of these samples, STRUCTURE was run using 62 reference individuals and the USEPOPINFO, POPFLAG, and PRCOMP settings. Reference individuals were chosen based on a Q-value > 0.94 when using the full GBS SNP dataset (14-24 per lineage). Other parameters: admixture model, K=3, correlated allele frequencies, migration prior 0.05, and burn-in of 20,000 followed by 150,000 MCMC iterations. A subset of 751 samples were also genotyped using a panel of 35 SNPs that had previously been developed for differentiating FLB and NLB, where ancestry proportions were determined by counting the number of FLB or NLB alleles using a custom R script (C. Li et al., 2015; Zhao et al., 2018).

### Genotype simulation and assignment

To evaluate the accuracy of our ancestry assignment using the 73 SNP panel and STRUCTURE, we employed a simulation approach developed by (Vähä & Primmer, 2006). Individuals used as the pure reference populations in our STRUCTURE runs (14 FLM, 24 DLB, 24 NLB) were used to simulate genotypes from random mating and hybridization with the hybridize R function in adegenet. One hundred genotypes were simulated for each pure lineage, 50 for each type of F1, 150 for each type of F2 through F4 generations assuming neutral admixture, and 300 triple hybrids (considered the product of crossing an F1 from a pair of species with a pure individual of the third species or a backcross between a triple hybrid and a pure species). These simulated genotypes were analysed using STRUCTURE with the same settings and reference genotypes. The performance of STRUCTURE for hybrid and pure individuals was evaluated based on efficiency (number of individuals in a simulated group that are correctly assigned), accuracy (proportion of an identified group that truly belongs to that group) and performance (efficiency multiplied by accuracy). Finally, the optimal threshold values of Q to assign individuals to the different genotypic categories was determined.

### Geographical patterns of Florida Bass introgression

To determine if patterns of FLB genetic ancestry could be explained by by both geographical variables and stocking history, we first calculated the great circle distance in km between our reference FLB population (St. Johns River, FL) and all other sites sampled with our SNP panel using the R package geosphere v1.5-10 (Hijmans et al., 2016). We then modeled the proportion of FLB ancestry for all 881 individuals as the interaction between distance from St. Johns River, FL and categorical stocking history, using a fractional logistic regression model implemented in base R stats [glm(FLB ancestry ~ distance * stocking, family = “quasibinomial”)]. Based on published literature and discussions with biologists at state agencies, sites were designated as either ‘River’ or ‘Reservoir’, where reservoirs were bodies of water with known stocking history and limited connectivity to other bodies of water (Alford & Jackson, 2009; Bunch et al., 2017; Hargrove et al., 2019). Ancestry proportions were those calculated using the 73 SNP diagnostic panel described below. This model was compared with a model based only on distance [glm(FLB ancestry ~ distance, family = “quasibinomial”)], by comparing the reduction in deviance with a chi-squared test.

## Results

### GBS Assembly

A total of 42,390 biallelic SNPs across 102,685 GBS loci were genotyped in greater than 50% of 144 largemouth bass individuals and at least three of the four groups (NLB, ILB, FLB, DLB). Mean length of GBS loci was 91 bp (sd=0.02). Further filtering by missing data < 15%, HWE, MAF >5%, and thinning by LD reduced the dataset to 8,582 SNPs. Average read depth across SNPs per individual ranged from 3.3 to 29.2 (mean = 11.2 ± 4.6).

### Population Genetics

A PCA of 144 individuals using 8,582 SNPs clearly differentiated the three lineages across PCs 1 and 2, with PC1 representing 30% of SNP variation and PC2 representing 11% variation (Figure 2). We calculated population genetic summary statistics on these 8,582 SNPs, with samples separated by either lineage group or sampling site. As expected, intergrade samples had the highest observed genome-wide heterozygosity (0.31), followed by DLB (0.16). NLB had the highest levels of inbreeding (0.105) (Table 1). Overall F_ST_ when separated by lineage group was 0.426. Pairwise F_ST_ between lineages was highest between FLB–NLB (0.749) and lowest between DLB–NLB (0.459). Fixed SNPs were detected between each pair of lineages, with the fewest between DLB–NLB (46 unlinked SNPs) and the most between FLB–NLB (1,807) (Table 2). There were 532 unlinked SNPs that were fixed between FLB–NLB but not between DLB–FLB, suggesting possible introgression between DLB–FLB. When comparing summary stats between sampling sites, Sugar Lake, MN had the lowest observed heterozygosity, followed by FLB samples from a captive hatchery population (Supp. Table 1). Pairwise F_ST_ between sites within a lineage group were generally low (mean pairwise F_ST_ 0.051 to 0.124), with the highest within-lineage F_ST_ observed between Sugar Lake, MN and Hatchery NLB (0.239) (Table 1).

**Figure 2.**
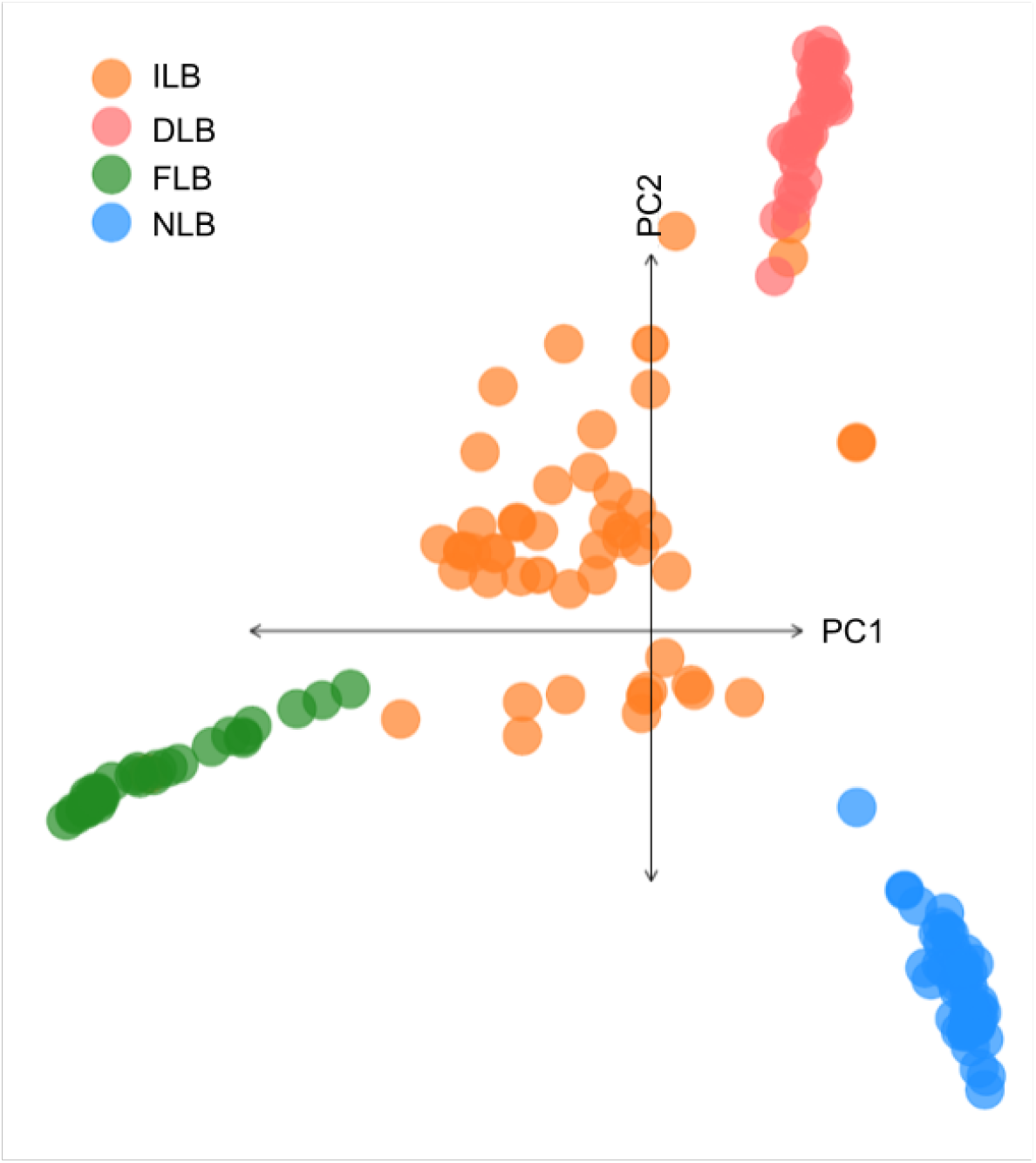
Principal component analysis plot of PC1 and PC2 for 144 *Micropterus* individuals based on 8,582 GBS SNPs. Individuals are colored by lineage (DLB=Delta Largemouth Bass, NLB=northern Largemouth Bass, FLB=Florida Bass, ILB=intergrade largemouth bass).

**Table 1.** Population genetics of largemouth bass lineages.

| Group | H<sub>o</sub> | H<sub>e</sub> | Avg. Pairwise F<sub>ST</sub> (Min-Max) | F<sub>IS</sub> |
| --- | --- | --- | --- | --- |
| ILB | 0.31 | 0.32 | 0.051 (0.0134-0.083) | 0.032 (0.027-0.036) |
| DLB | 0.16 | 0.17 | 0.031 (0.008-0.05) | 0.070 (0.063-0.077) |
| FLB | 0.11 | 0.13 | 0.09 (0.065-0.11) | 0.079 (0.069-0.089) |
| NLB | 0.12 | 0.14 | 0.124 (0.037-0.239) | 0.105 (0.097-0.112) |

Population genetic summary statistics based on 8,582 SNPs for 144 individuals across the three largemouth bass lineages and their intergrades (DBS=Delta Largemouth bass, FLB=Florida Bass, NLB=northern Largemouth bass, ILB=intergrades). H<sub>o</sub>: observed heterozygosity averaged across loci; H<sub>e</sub>: expected heterozygosity averaged across loci; F<sub>IS</sub>: Wright's F-statistics averaged across loci (Nei & Chesser, 1983).

**Table 2.** Genetic differentiation of largemouth bass lineages.

|  | ILB | DLB | FLB | NLB |
| --- | --- | --- | --- | --- |
| DLB | 0.203 | 0 | 1,275 (15%) | 46 (0.5%) |
| FLB | 0.329 | 0.677 | 0 | 1,807 (21%) |
| NLB | 0.347 | 0.459 | 0.749 | 0 |

Lower triangle: pairwise F<sub>ST</sub> between lineages of largemouth bass and intergrades using 8,582 GBS SNPs. Upper triangle: number and percentage of unlinked, “fixed” (> 0.98 allele frequency) SNPs between lineages.

### Phylogeny and inferred ancestry

Maximum likelihood phylogenetic inference using 18,492 GBS SNPs supported DLB as a distinct lineage that is sister to NLB, with 100% support from both the SH-like approximate likelihood ratio test (SH-aLRT) and ultrafast bootstrapping (Figure 3). When samples from intergrade populations were included, SH-aLRT support for DLB as a unique lineage was 99.9% and ultrafast bootstrap support was 96% (Supp. Fig. 1). For the first 598 bp of the ND2 mitochondrial gene, there were two Delta haplotype, ten FLB haplotypes, and ten NLB haplotypes (Figure 4). There were two fixed ND2 SNPs between DLB-NLB, 21 between DLB-FLB, 19 between FLB-NLB. Intergrade samples, which had STRUCTURE-inferred ancestry proportions of at least 6% in two or more lineages, were distributed across both the GBS phylogenetic tree and across mitochondrial haplotypes (Supp. Fig. 1, Figure 4). Individuals from some sampling sites, including Sugar Lake, MN and Demopolis Lake, AL, sequenced well on the MassARRAY platform but did not amplify or sequence well with available ND2 primers (Kocher et al., 1995), suggesting a polymorphism in the ND2 primer recognition sites. Some intergrade individuals appeared to be heterozygous at ND2 sites that were diagnostic between lineages, which may be a sign of heteroplasmy (Piganeau et al., 2004) or sequencing of homologous regions in the nuclear genome (nuMTs) (Simone et al., 2011). This included two individuals from West Point Reservoir, GA, one from Lake Eufaula, AL, one from Hatchie River, TN, and one from Sutton Lake, NC. All of these sites have known recent stocking history.

**Figure 3.**
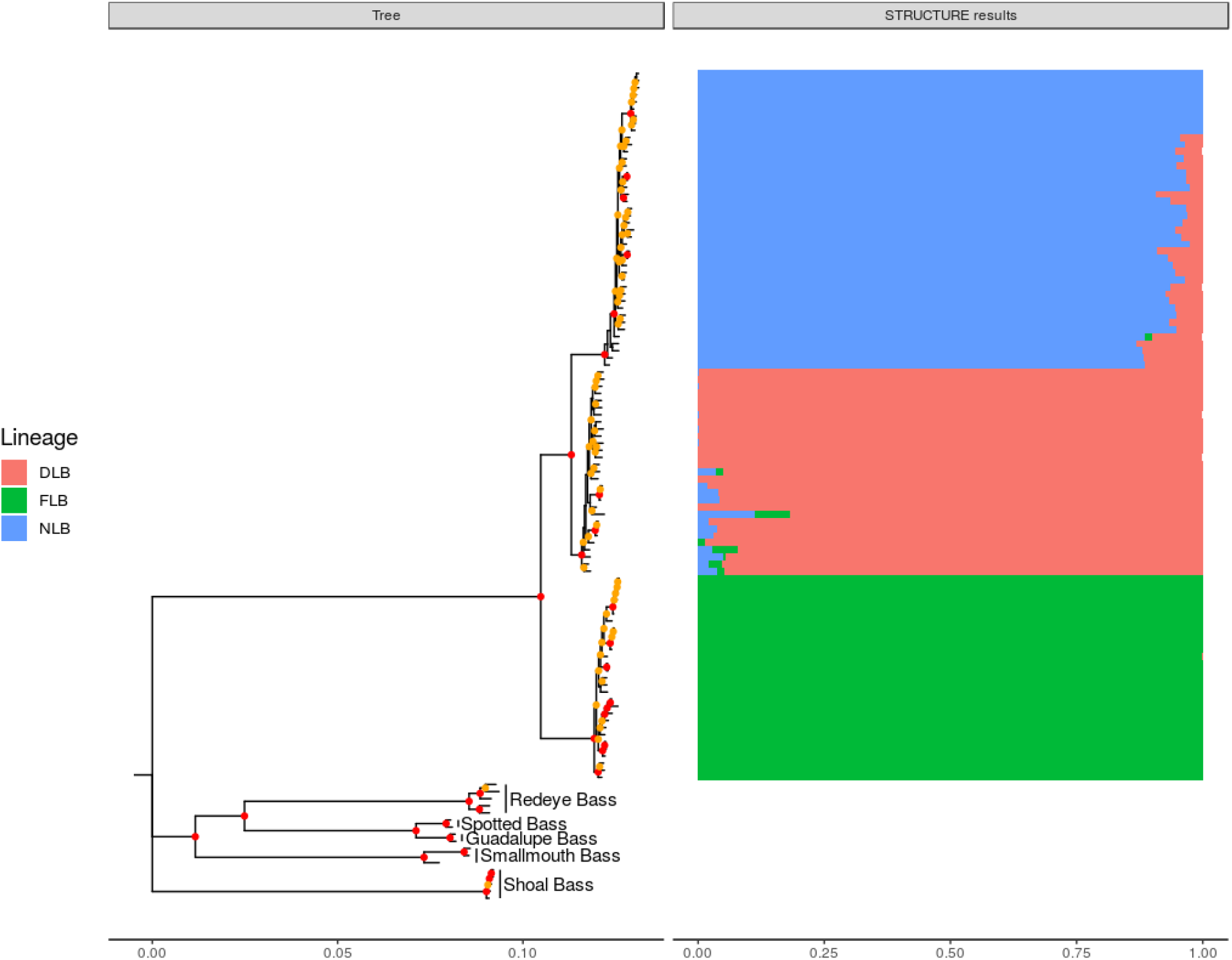
Maximum likelihood phylogeny constructed using 18,538 concatenated GBS SNPs and IQ-TREE (1000 ultrafast bootstrap replicates). Nodes in red indicate 100% ultrafast bootstrap support, nodes in yellow indicate 50-99% ultrafast bootstrap support. STRUCTURE plots (K=3) are included and show the ancestry membership proportions (Q-values) for the three largemouth bass lineages as inferred using 8,582 SNPs (DLB=Delta Largemouth Bass, NLB=northern Largemouth Bass, FLB=Florida Bass).

**Figure 4.**
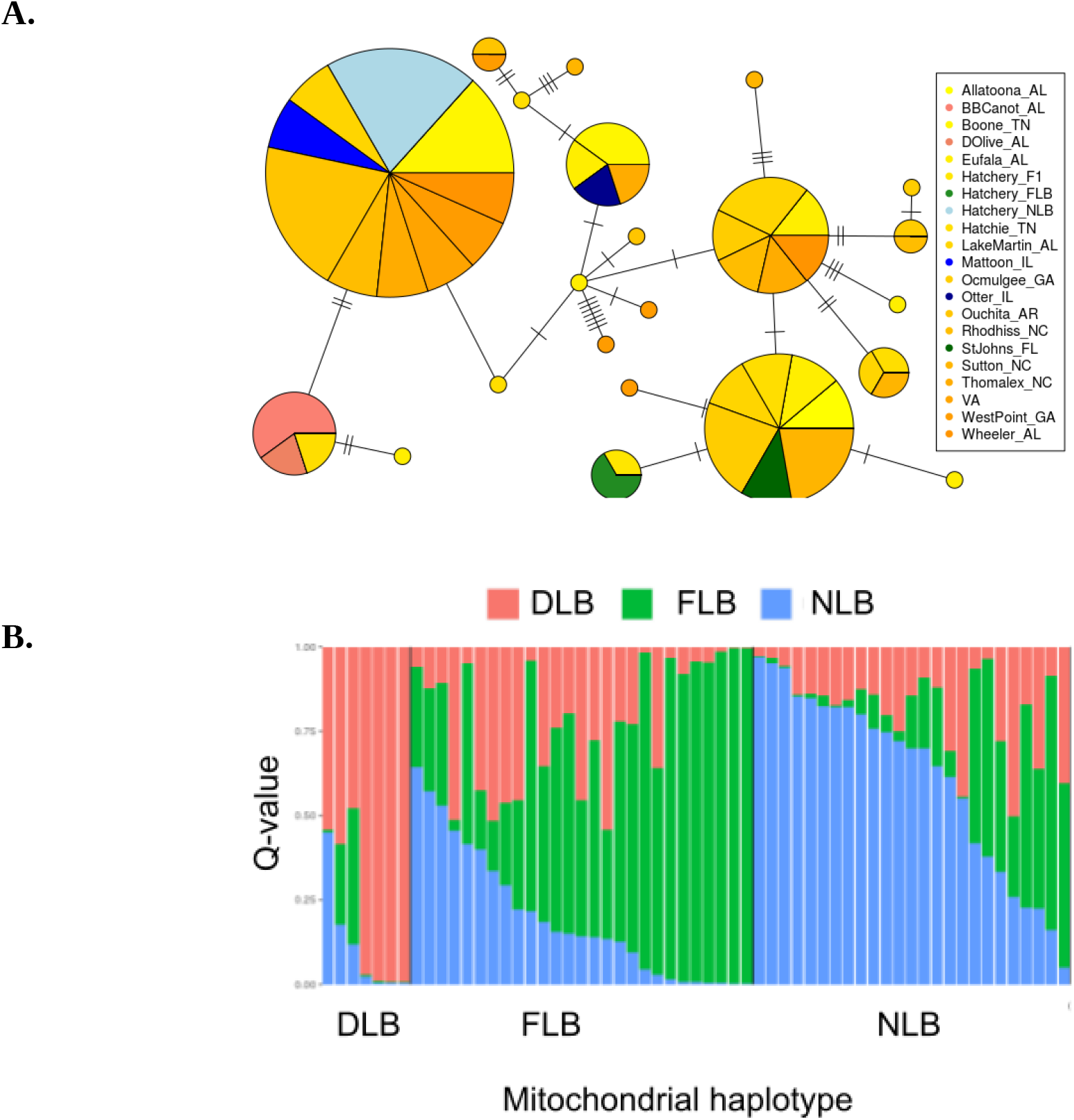
**A)** Minimum spanning network of mitochondrial haplogroups from 62 individuals, based on the first 529 bp of the ND2 gene. Circle size is proportional to haplotype frequency and colors represent sampling location. **B)** STRUCTURE (K=3) results based on 73 diagnostic SNPs, ordered by mitochondrial haplotype group (DLB=Delta Largemouth Bass, FLB=Florida Bass, NLB= northern Largemouth Bass).

### Panel Development

From the 127 diagnostic SNPs retained for panel development, the MassARRAY Assay Design software developed primers for 91 SNPs across two multiplexed panels (52 and 39). Detailed information on SNP panels, including SNP ID, genome position, and primer sequences (forward, reverse, and extension), are listed in Supp. File 3. After removing SNPs that sequenced poorly or were invariant, 73 SNPs were retained and run on 881 individuals. STRUCTURE results on the 144 GBS individuals using 8,582 GBS SNPs, 2,809 diagnostic GBS SNPs, and the 73 panel SNPs were highly concordant (Fig. 5). We also examined the consistency of MassARRAY genotype calls among 70 technical replicates and found 98.4% of genotypes matched across multiple runs. Analysis of simulated genotypes indicated that a threshold value in the admixture coefficient of Q ≥ 0.94 resulted in adequate separation between pure species and hybrid individuals (Table 3). The genetic assignment of simulated pure individuals from the three lineages showed high levels of performance (98%–100%). For hybrid classes involving two lineages and fewer than four generations of backcrossing, performance values ranged from 99%–93%. While individuals were classified as NLB–FLB hybrids with 100% accuracy, 6.5% of the simulated NLB–FLB hybrids were misclassified as triple hybrids, showing a small observed DLB Q-value (< 0.1). Triple hybrids had > 90% performance. When fourth generation backcrossed hybrids (Bx4) were included, efficiency decreased to 82.8%-92.4% for hybrids and accuracy decreased to 58%-78.7% for pure species, as many Bx4 hybrids were assigned as pure (Supp. Table 2).

**Figure 5.**
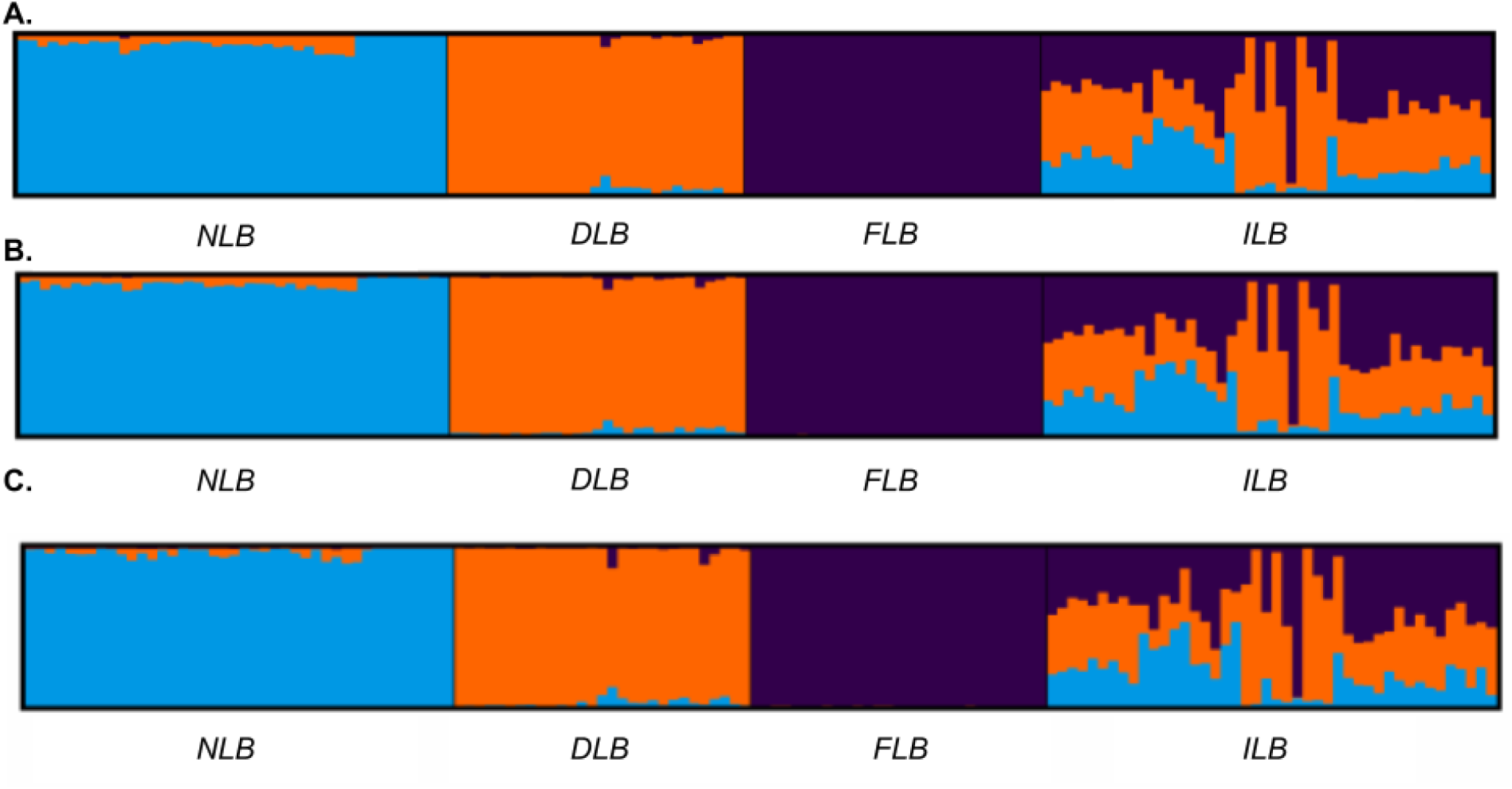
STRUCTURE results (K=3) from three GBS SNP subsets (K=3). A) 8,582 SNPs, B) 2,809 diagnostic SNPs, and C) 73 SNPs chosen for SNP panel. NLB=northern Largemouth Bass, DLB=Delta Largemouth Bass, FLB=Florida Bass, ILB=intergrade largemouth bass.

**Table 3:** Evaluation of 73 SNP panel on simulated genotypes.

| Simulated/<br>Assigned | DLB | FLB | NLB | DLB-FLB | DLB-NLB | NLB-FLB | Triple<br>Hybrid | Total<br>Simulated |
| --- | --- | --- | --- | --- | --- | --- | --- | --- |
| DLB | **100** |  |  |  |  |  |  | 100 |
| FLB |  | **100** |  |  |  |  |  | 100 |
| NLB |  |  | **100** |  |  |  |  | 100 |
| DLB-FLB F1 |  |  |  | **50** |  |  |  | 50 |
| DLB-FLB<br>F2:F4 |  |  |  | **149** |  |  | 1 | 150 |
| DLB-FLB BX | 2 |  |  | **198** |  |  |  | 200 |
| DLB-NLB F1 |  |  |  |  | **50** |  |  | 50 |
| DLB-NLB<br>F2:F4 |  |  |  |  | **150** |  |  | 150 |
| DLB-NLB BX |  |  | 1 |  | **199** |  |  | 200 |
| NLB-FLB F1 |  |  |  |  |  | **47** | 3 | 50 |
| NBB-FLB<br>F2:F4 |  |  |  |  |  | **131** | 19 | 150 |
| NLB-FLB BX |  |  |  | 1 | 2 | **193** | 4 | 200 |
| Triple Hybrid |  |  |  | 4 |  |  | **296** | 300 |
| Total<br>Assigned | 102 | 100 | 101 | 402 | 401 | 371 | 323 | 1800 |
| **Efficiency** | 100.00% | 100.00% | 100.00% | 99.25% | 99.75% | 92.75% | 98.67% |  |
| **Accuracy** | 98.04% | 100.00% | 99.01% | 98.76% | 99.50% | 100.00% | 91.64% |  |
| **Performance** | 98.04% | 100.00% | 99.01% | 98.02% | 99.25% | 92.75% | 90.42% |  |

### Geographical patterns of ancestry and hybridization

Population structure analysis of 52 sites using the 73 SNP panel identified populations with a majority of pure DLB individuals extending north of the Mobile Delta, encompassing all tested populations below the Fall Line, including those in remote areas with no history of stocking (e.g., Sipsey River; Fig. 1). Pure FLB individuals were found far outside of the currently accepted Florida Bass native range, extending into Georgia south of the Fall Line and up the Atlantic coastal plain into North Carolina, while intergrade populations between FLB and NLB were also found outside of the intergrade zone as described by Bailey and Hubbs (1949). Twenty-six intergrade populations with > 6% mean ancestry from all three lineages were identified, primarily in Alabama, Georgia, Tennessee, and Louisiana. Compared to previous estimates of FLB–NLB ancestry using a 35 SNP panel, this 73 SNP panel appears to more accurately represent FLB ancestry in sites with high DLB ancestry. For example, while the 35 SNP panel found that populations from the Mobile Delta were 27–30% FLB and 70–73% NLB, the 73 SNP panel identifies them as 95–98% DLB (Supp. Fig. 3, Supp. File 2).

To evaluate the relative roles of geography and stocking history for determining FLB genetic ancestry, we implemented fractional logistic regression models based on the distance of a site from our reference FLB population and the stocking history of a site (coded as either River or Reservoir). The proportion of FLB ancestry in an individual was best described by a model that included stocking: FLB ancestry ~ distance * stocking (Chi-squared test of reduction in deviance, p-value = 7.43e-12), where separate geographic clines were fit for River and Reservoir sites (Figure 6). Based on values predicted by the model for Reservoir sites, stocking has increased the overall geographic extent of FLB intergrades (FLB ancestry > 0.06) by 436 km on average. However, the model also indicates that individuals with pure FLB ancestry observed in Georgia below the Fall Line are likely not the result of anthropogenic movement. Both the model and our observations in unstocked River sites support a larger natural distribution of pure FLBs and intergrades than is currently accepted (A. T. Taylor et al., 2019).

**Figure 6.**
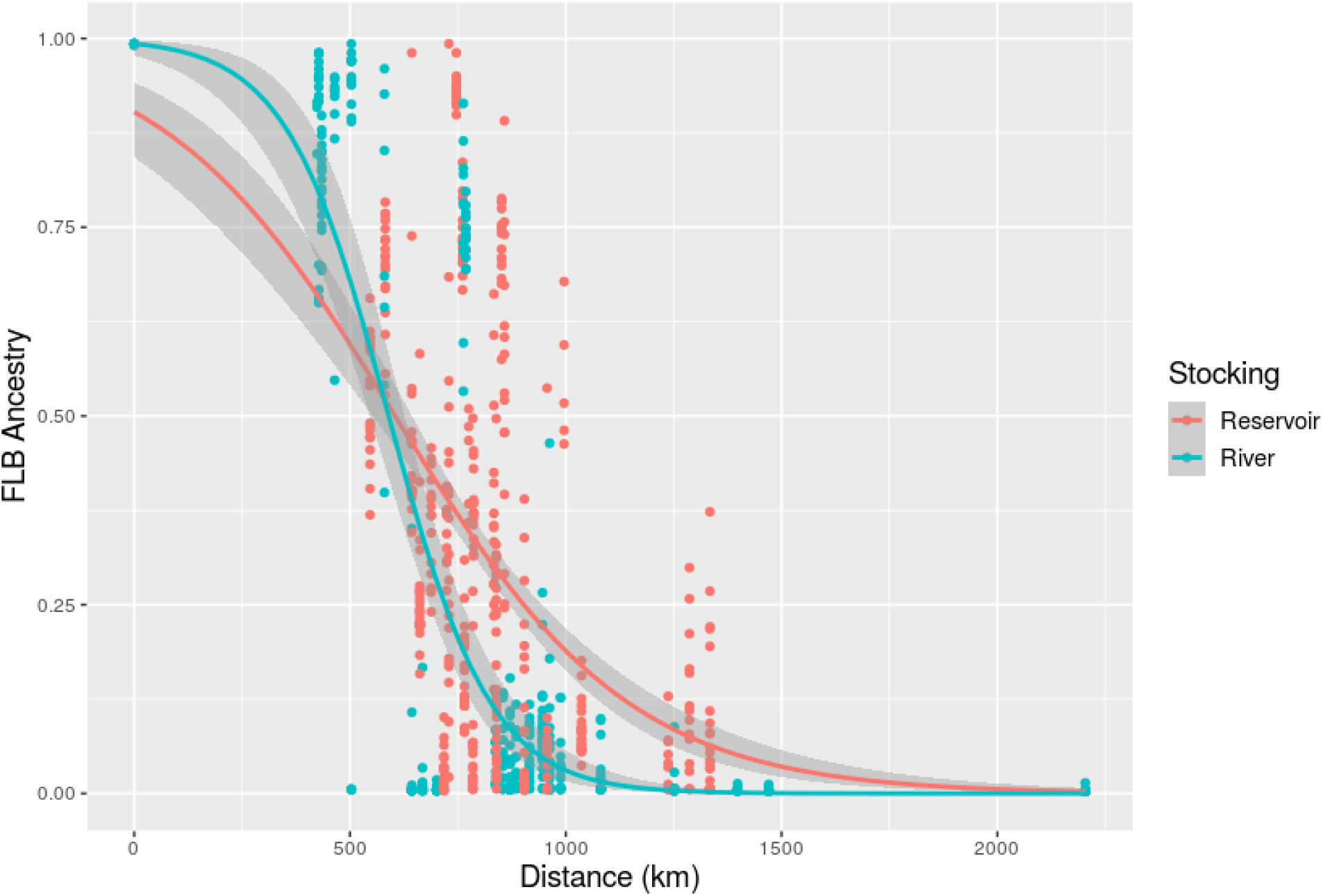
Scatterplot of *M. floridanus* (FLB) ancestry proportions for 881 individuals, plotted by great circle distance from a reference pure FLB site in St. Johns River, FL. FLB ancestry proportions are based on STRUCTURE results (K=3) using a panel of 73 diagnostic SNPs. Individuals are colored by stocking history (River= no significant FLB stocking, Reservoir=known stocking history). Lines show the fit of a fractional logistic regression model (FLB ~ distance * stocking).

## Discussion

Using a combination of 8,582 SNPs and targeted mitochondrial gene sequencing, we identified a complex hybrid zone involving three distinct lineages of largemouth bass (*M. floridanus* and two from *M. salmoides*), including the first thorough genetic characterization of a unique lineage of *M. salmoides* radiating out from the Mobile-Tensaw Delta. To assess the geographic extent of hybridization, we developed a suite of 73 SNPs that can accurately classify advanced-generation hybrids between all three lineages. This SNP panel will be a valuable tool for fisheries management and conservation of these economically and ecologically important species, while also facilitating future investigations of introgression and species boundaries. While stocking of Florida Bass in reservoirs for recreational fishing has contributed in part to observed hybridization patterns, the natural geographic extent of both pure *M. floridanus* and introgressed populations are nonetheless much larger than currently accepted.

### Distinct Mobile Delta lineage of Largemouth Bass

Support from both nuclear and mitochondrial data indicate that Largemouth Bass from the Mobile-Tensaw Delta (DLB) are an evolutionarily diverged lineage of *M. salmoides*. Genetic differentiation between DLB and northern Largemouth Bass (NLB) was high (F_ST_ = 0.459), with 46 unlinked nuclear SNPs fixed between the two groups and two fixed SNPs in the first 598 bp of the mitochondrial ND2 gene. While significant, this differentiation is still much lower than that observed between Florida Bass (FLB) and NLB (F_ST_=0.749, 1,807 fixed unlinked nuclear SNPs, 19 fixed ND2 SNPs). These values are higher than those observed using GBS SNPs for different species of *Squalius* chubs (F_ST_=0.126–0.414) and comparable to F_ST_ between diverged lineages of walleye (0.805) (Mendes et al., 2021; Zhao et al., 2020). Our results contribute to a growing scientific consensus around the validity of *M. floridanus* as a distinct species (Kassler et al., 2002; Near et al., 2003; Near & Kim, 2021). While this study cannot address whether Delta bass should be considered a new species or subspecies of *M. salmoides*, it does indicate that morphological evaluations aided by genetic assays are warranted.

Maximum likelihood phylogenetic analysis of nuclear loci and a haplotype network analysis of a mitochondrial locus suggest that DLB–NLB diverged more recently than NLB–FLB. Fossil-calibrated phylogenies of *Micropterus* using either two mitochondrial genes or 16 nuclear genes dated the divergence of NLB–FLB at the end of the Pliocene (~3.3 Mya) (Near et al., 2003; Near & Kim, 2021), suggesting that a NLB population in the Mobile Delta may have diverged later during the Pleistocene, when glaciation and sea level changes resulted in fragmented coastal freshwater habitat (Bermingham & Avise, 1986). Once the population was reconnected with the mainland, we hypothesize these Delta Largemouth Bass migrated out of the Delta and met with inland *M. salmoides* and *M. floridanus*. The Mobile Delta region is also home to genetically diverged populations of walleye and yellow perch, although the estimated divergence time between these relict populations and their northern counterparts range from 1.7 Mya for walleye to 170,000 ya for perch (Stepien et al., 2015), suggesting that the timing of population divergence in this geographic locale can vary across fish taxa.

Genome-wide genetic diversity was higher in pure DLB populations compared to pure populations of FLB or NLB (Table 1, Supp. Table 1). As we only had two to four wild populations of each lineage, this may reflect our limited geographical sampling. This result contrasts with walleye and perch diversity, where their Mobile Delta populations had lower genetic diversity than those in the north (Stepien et al., 2015). Elevated genetic diversity is encouraging, as it suggests that the DLB populations have not experienced dramatic population declines and bottlenecks (Nei et al., 1975). Historical or recent introgression with FLB is one potential explanation for higher genetic diversity in DLB populations. Genetic divergence between DLB–FLB is lower than between NLB–FLB (Table 2), supporting this hypothesis. Introgression between the two may have occurred naturally across the Florida panhandle, or due to unsuccessful stocking efforts that occurred in the Mobile Delta sporadically from 1988 to 2000 (Armstrong et al., 2000; Rainer, 2010). Sugar Lake, MN had the lowest genetic diversity out of all sampled sites, which may be due to a recent genetic bottleneck and range expansion after the last glacial maximum (Dyke & Prest, 1987). Sugar Lake is also the only NLB population with > 99% mean inferred NLB ancestry based on STRUCTURE. We hypothesize that the low levels of DLB ancestry (< 5%) observed in other “pure” NLB populations, such as those in Illinois, are likely due to ancestral polymorphisms. The potential genetic bottleneck experienced by Sugar Lake may have removed some of these shared variants between DLB and NLB from the population.

### Multi-lineage hybrid zone

The hybrid zone between NLB and FLB was first described by Bailey and Hubbs in 1949 using meristics, and found to extend only through Georgia and a small part of Alabama (Bailey & Hubbs, 1949). Although subsequent studies using larger sample sizes and biochemical or genetic assays have determined the geographic extent of hybridization to be much larger, these results are often discounted as merely the product of *M. floridanu*s recreational stocking outside of Florida. Using our diagnostic 73 SNP panel, we assayed 52 populations across the east and southeast, representing both reservoirs with known stocking history and unaltered river sites (Fig. 1). We found that stocking did not explain the presence of FLB intergrades observed in North Carolina, however stocking has likely increased the mean percent of FLB ancestry in those reservoirs. Unstocked river populations with many pure FLB fish were observed in Georgia south of the Fall Line, which is also in contrast to the currently accepted native range of *M. floridanus*. These results strongly support a redrawing of native ranges for both the intergrade zone and *M. floridanus*. To fully describe the extent of introgression in these lineages, more geographic sampling with the 73 SNP panel is needed, particularly in Mississippi and South Carolina.

One surprising result of our study was the high number of hybrids detected with ancestry contributions from all three lineages (367, 41% of samples analyzed with 73 SNP panel). As our panel was shown to have an 8.4% false discovery rate (Table 3), we can estimate that at least 336 of these triple hybrids are real. Populations with high proportions of triple hybrids (> 20%) were primarily in the Atchafalaya Basin in Louisiana, reservoirs of Alabama and Georgia, and two sites in Tennessee. While the low FLB ancestry observed in Louisiana and Tennessee are likely due to stocking (Hargrove et al., 2019), we hypothesize that the triple hybrids in Georgia and Alabama are a result of secondary contact among the three lineages after allopatric divergence. Hybrids between three diverged lineages are more commonly detected in plants, but have been observed in some fishes such as cichlids, suckers, and darters (Keck & Near, 2009; McDonald et al., 2008; Nevado et al., 2011).

Whether the result of natural introgression, anthropogenic introductions, or a combination of the two, the large hybrid zone observed between these three diverged lineages provides an excellent opportunity to study the speciation process (Barton & Hewitt, 1985; Gompert et al., 2017). When hybrid zones act as semi-permeable barriers to gene flow between parental species, patterns of introgression can vary across the genome (Harrison & Larson, 2014). Differential introgression among loci can be due to stochastic processes (i.e., drift) or selection associated with genotype-by-environment interactions or reproductive isolation (Fitzpatrick et al., 2009; Gompert et al., 2012; S. A. Taylor et al., 2015). With additional genomic sequencing and a careful geographic sampling design aided by the diagnostic panel produced in this study, largemouth bass can be developed into a new case study for genomic patterns of introgression when multiple lineages are involved.

### SNP assay for hybridization and species identification

Characterizing genomic contributions of hybrids is critical for understanding hybrid zone dynamics and informing fisheries management (Fitzpatrick et al., 2015). Empirical and simulation-based studies have determined that at least 50 ancestry-informative genetic markers may be required for accurately classifying F2 hybrids and advanced-generation backcrossed individuals (Fitzpatrick, 2012; Malde et al., 2017). Previous work has resulted in panels of 18 microsatellites or 25-38 SNPs for assessing integrity and hybridization between *M. salmoides/M. floridanus*, however these panels did not include samples from the Mobile-Tensaw Delta during development and therefore may not accurately describe patterns of introgression (C. Li et al., 2015; Seyoum et al., 2013; Zhao et al., 2018). When comparing results to our 73 SNP panel, we find that *M. floridanus* ancestry inferred with a previous 35 SNP FLB–NLB assay is often inflated in sites with high DLB ancestry. For example, populations of pure (>94%) DLB ancestry in the Mobile Delta were shown to have >25% FLB ancestry when using the previous assay (Supp. Fig. 3, Supp. File 2).

Designing a diagnostic SNP panel for more than two species is a challenge, as fixed triallelic SNPs are rare and therefore only biallelic SNPs are retained. Unlike a previous NLB–FLB SNP assay, we are not able to simply count the number of FLB and NLB alleles to determine hybrid ancestry proportions. Instead, we developed a rigorous approach for screening diagnostic loci by leveraging the program STRUCTURE and reference genotypes for accurate estimation of ancestry. Our panel of 73 SNPs captures the same ancestry proportions as the full set of 8,582 GBS SNPs for all individuals within Q-values of 0.1. By evaluating our panel on simulated hybrid genotypes, we also determined that a Q-value cutoff of ≥ 0.94 was accurate for classifying whether an individual was likely derived from only one lineage. Many other studies that use STRUCTURE to identify pure individuals of a species use an arbitrary Q-value cutoff, such as ≥0.95 (Lutz-Carrillo et al., 2006; Thongda et al., 2020) or ≥ 0.90 (Dakin et al., 2015). When utilizing a reduced number of diagnostic markers in STRUCTURE, simulations should be applied to ascertain performance at various Q-value thresholds.

### Management implications for largemouth bass

As highly popular sportfish species, management of NLB and FLB involves balancing angler-desirable traits (e.g., size, catchability), stable fisheries, and biodiversity concerns (Young et al., 2006). The phenotypic and demographic impacts of anthropogenic hybridization between the two species have been studied extensively, with some finding negative fitness consequences due to outbreeding depression (Cooke et al., 2001; Cooke & Philipp, 2006; Philipp & Claussen, 1995) and others seeing a potential to enhance managed fisheries (Maceina et al., 1988; Maceina & Murphy, 1992). Many of these early studies failed to take genetic ancestry into account or evaluated hybrids between geographically disparate sites, making it difficult to extrapolate results to naturally occurring intergrades. The discovery of a genetically distinct lineage of Largemouth Bass that may differ in fishery-relevant traits (e.g., smaller size, slower growth rate, and environmental tolerances) further strengthens the need for accurate genetic tools to determine purity and hybridization. To better inform stocking success, future performance assays should assess genetic ancestry and leverage natural intergrades to provide a complete picture of fitness variation across largemouth bass lineages.

Current diagnostic markers in largemouth bass include a set of 35 SNPs designed for the MassARRAY Sequenom (C. Li et al., 2015; Zhao et al., 2018) and a panel of 18 microsatellites (Barthel et al., 2010; Seyoum et al., 2013). While we have not evaluated our new panel against these microsatellites, they likely face the same issues as the 35 SNP panel for mischaracterizing ancestry. Given the numerous populations with Delta bass ancestry observed in the southeast, genetic markers used by state agencies and evolutionary biologists need to be updated to better characterize the geographic extent of pure and hybrid populations of all three lineages. For example, some sites in Alabama and Georgia that were thought to have roughly the same FLB–NLB ancestry are now shown to differ considerably in their proportions of DLB ancestry. The continued movement and stocking of largemouth bass by state agencies and commercial hatcheries makes this a pressing issue. Genotyping of 5 ‘pure’ northern Largemouth Bass from a popular commercial hatchery found DLB ancestry ranging from 0.02–0.17. Without understanding the potential genotype-by-environment interactions of these hybrids, DLB alleles may be spread far outside of their native range to detrimental outcomes.

The observation of individuals with DLB mitochondria and ancestry extending outside of the Mobile Delta raises the question of whether phenotypic variation observed in fish from the Delta are due to adaptation or phenotypic plasticity from the environment. While Largemouth Bass populations of the Mobile Delta are currently stable, they may be at particular risk from climate change due to unique environmental challenges in coastal habitats, such as low dissolved oxygen or storm-mediated shifts in salinity. Microcosm experiments and functional genomics with both inland and coastal Delta bass can help elucidate whether these fish have adapted or acclimated to the estuarine environment.

## Data Accessibility

Raw demultiplexed Illumina DNA sequences were previously published (Thongda, et al., 2020) and are available on NCBI SRA PRJNA417468. The draft *M. floridanus* genome assembly was previously published (Zhao, et al., 2018), and is available on DDBJ/ENA/GenBank under the accession NRCI01000000. Genomic data (all filtered SNPs, all diagnostic SNPs, and MassARRAY genotypes for the 73 SNP panel) will be available on Dryad at time of publication. ND2 gene sequences will be available on GenBank at time of publication. All code will be available on the author’s Github, and can be made available to reviewers by request.

## Note

We adopt the species designation of Florida Bass (*Micropterus floridanus*) as supported by the genetic work of Kassler et al., (2002) and Near et al., (2003), although the American Fisheries Society’s Names of Fishes Committee continues to use the subspecies designation Florida Largemouth Bass (*Micropterus salmoides floridanus*) (Page et al., 2013).

## Acknowledgements

This work was supported by funding from the Southeastern Fish Genetics Cooperative (E. Peatman, director), particularly with support from Alabama Department of Conservation and Natural Resources. We thank all individuals and state agencies who provided samples or assisted with sample collections, including Alabama Dept of Conservation and Natural Resources, Georgia Dept of Natural Resources, Tennessee Wildlife Resources Agency, Louisiana Dept of Natural Resources, North Carolina Wildlife Resources Commission, Delaware Dept of Natural Resources and Environmental Control, Virginia Dept of Wildlife Resources, and Maryland Dept of Natural Resources.

## Supplementary File 1

**Supp. Table 1:** Sample information for GBS individuals. Population genetic summary statistics are provided for GBS sampling sites with at least 5 individuals using 8,582 SNPs. H_o_: observed heterozygosity averaged across loci; H_e_: expected heterozygosity averaged across loci; F_IS_: Wright’s F-statistics averaged across loci (Nei & Chesser, 1983).

| Population | Group | H<sub>o</sub> | H<sub>e</sub> | F<sub>IS</sub> | N | Latitude | Longitude |
| --- | --- | --- | --- | --- | --- | --- | --- |
| Lake Allatoona, GA | ILB | 0.277 | 0.295 | 0.062 (0.054-0.072) | 9 | 34.1331 | -84.6244 |
| Lake Guntersville, AL | ILB | 0.346 | 0.324 | -0.068 (-0.075--0.060) | 10 | 34.5523 | -86.1192 |
| Lay Lake, AL | ILB | 0.252 | 0.286 | 0.118 (0.110-0.127) | 9 | 33.1166 | -86.482 |
| Lake Eufaula, AL | ILB | 0.276 | 0.291 | 0.051 (0.040-0.062) | 5 | 31.91168 | -85.15915 |
| Lake Harding, AL | ILB | 0.315 | 0.310 | -0.016 (-0.028 -0.004) | 5 | 32.6911 | -85.1114 |
| Rocky Mountain PFA, GA | ILB | 0.287 | 0.305 | 0.059 (0.0475-0.072) | 5 | 34.34236 | -85.32919 |
| Big Bayou Canot, AL | DLB | 0.145 | 0.150 | 0.032 (0.015-0.048) | 6 | 30.83575 | -87.98424 |
| D'Olive Bay, AL | DLB | 0.140 | 0.145 | 0.036 (0.021-0.050) | 8 | 30.65620 | -87.41795 |
| Sipsey River, AL | DLB | 0.167 | 0.171 | 0.020 (0.011-0.031) | 13 | 33.91454 | -87.68724 |
| St. Johns River, FL | FLB | 0.122 | 0.122 | -0.0004 (-0.015- 0.013) | 17 | 28.24763 | -81.38371 |
| Hatchery FLB | FLB | 0.093 | 0.105 | 0.103 (0.079-0.127) | 7 | NA | NA |
| FL Bass Conservation Center | FLB | 0.114 | 0.112 | -0.015 (-0.038 -0.007) | 5 | 28.50138 | -82.05196 |
| Lake Mattoon, IL | NLB | 0.139 | 0.136 | -0.02 (-0.032 - -0.007) | 10 | 39.34934 | -88.47698 |
| Hatchery NLB | NLB | 0.125 | 0.128 | 0.022 (0.006-0.039) | 6 | NA | NA |
| Otter Lake, IL | NLB | 0.139 | 0.143 | 0.031 (0.018-0.044) | 10 | 39.41576 | -89.90370 |
| Sugar Lake, MN | NLB | 0.077 | 0.080 | 0.031 | 9 | 45.31384 | -94.04487 |
|  |  |  |  | (0.011-0.050) |  |  |  |
| Lamar County Lake, AL | ILB | NA | NA | NA | 1 | 33.7769 | -88.2349 |
| Reeves Branch, Chattahoochee R., AL | ILB | NA | NA | NA | 5 | 31.88108 | -85.1875 |
| Reelfoot Reservoir, TN | ILB | NA | NA | NA | 4 | 36.40582 | -89.3809 |
| Tensaw Lake, AL | DLB | NA | NA | NA | 2 | 31.03156 | -87.90164 |

**Supp. Table 2:** Accuracy and efficiency of 73 SNP panel when simulated 4th generation backcrossed individuals are included. Number of simulated individuals (rows), which were assigned to one of three lineages or into a hybrid category (columns). Simulated individuals were assigned based on their STRUCTURE Q-values, with a threshold of Q ≥ 0.94 for pure individuals. Parameters of efficiency, accuracy and overall performance of the assignment method are given in percent. Individuals correctly assigned are in bold type. DLB: Delta Largemouth Bass (*M. salmoides*), NLB: northern Largemouth Bass (*M. salmoides*), FLB: Florida Bass (*M. floridanus*); BX: backcrosses.

| Simulated/<br>Assigned | DLB | FLB | NLB | DLB-FLB | DLB-NLB | NLB-FLB | Triple<br>Hybrid | Total<br>Simulated |
| --- | --- | --- | --- | --- | --- | --- | --- | --- |
| NB-FL BX4 |  | 27 | 26 |  | 3 | **43** | 1 | 100 |
| DB-NB BX4 | 19 | 18 |  |  | **63** |  |  | 100 |
| DB-FL BX4 | 28 | 27 |  | **45** |  |  |  | 100 |
| Total | 47 | 72 | 26 | 45 | 66 | 43 | 1 |  |
| Efficiency | 100% | 100% | 100% | 88.40% | 92.40% | 82.80% | 98.67% |  |
| Accuracy | 67.11% | 58.14% | 78.74% | 98.88% | 98.93% | 100.00% | 91.36% |  |
| Performance | 67.11% | 58.14% | 78.74% | 87.41% | 91.41% | 82.80% | 90.14% |  |

**Supp. Fig. 1.**
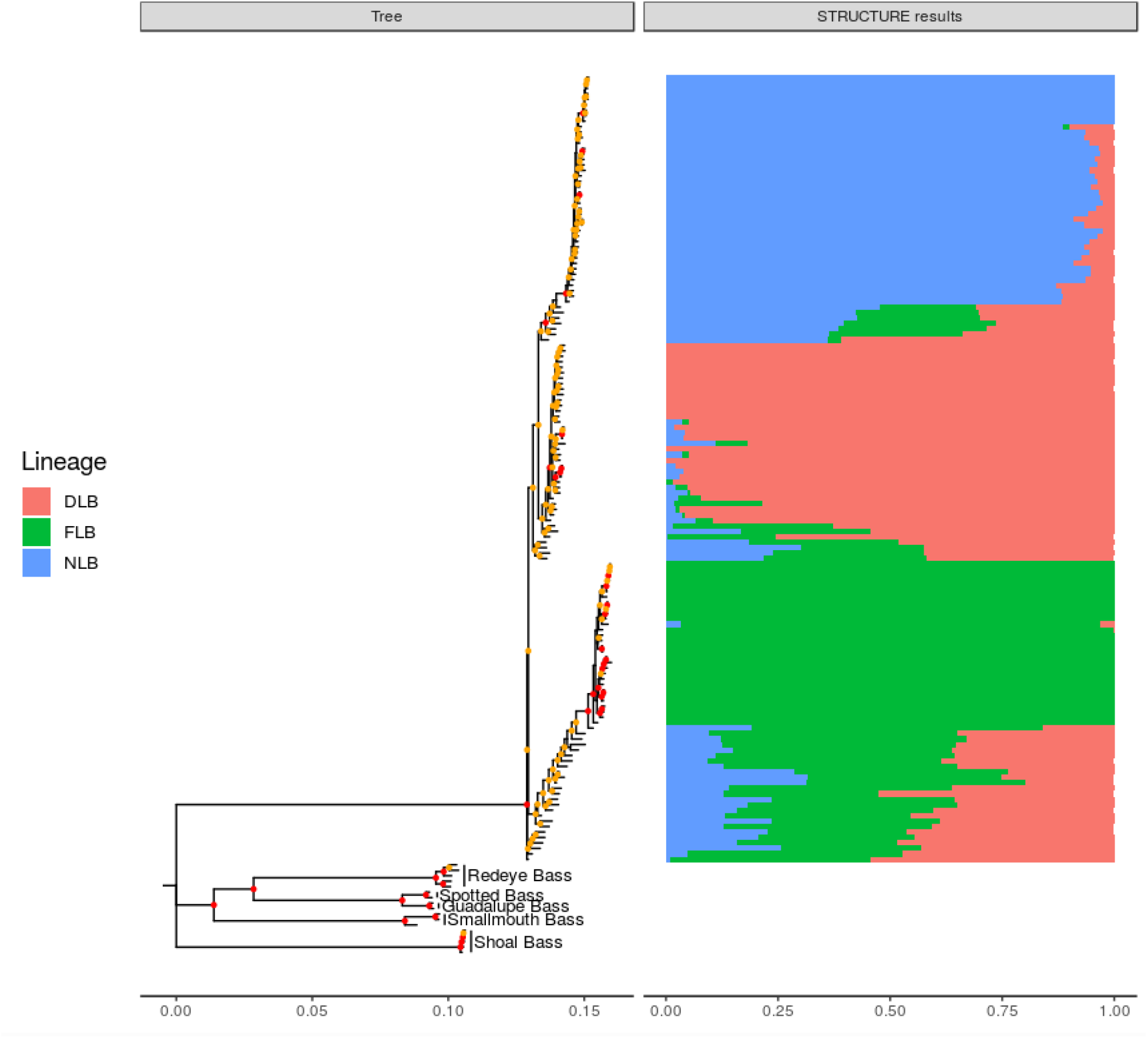
Maximum likelihood phylogeny constructed using 18,538 concatenated GBS SNPs and IQ-TREE (1000 ultrafast bootstrap replicates). Nodes in red indicate 100% ultrafast bootstrap support, nodes in yellow indicate 50-99% ultrafast bootstrap support. STRUCTURE plots (K=3) are included and show the ancestry membership proportions (Q-values) for the three largemouth bass lineages as inferred using 8,582 SNPs (DLB=Delta Largemouth Bass, NLB=northern Largemouth Bass, FLB=Florida Bass).

**Supp. Figure 2:**
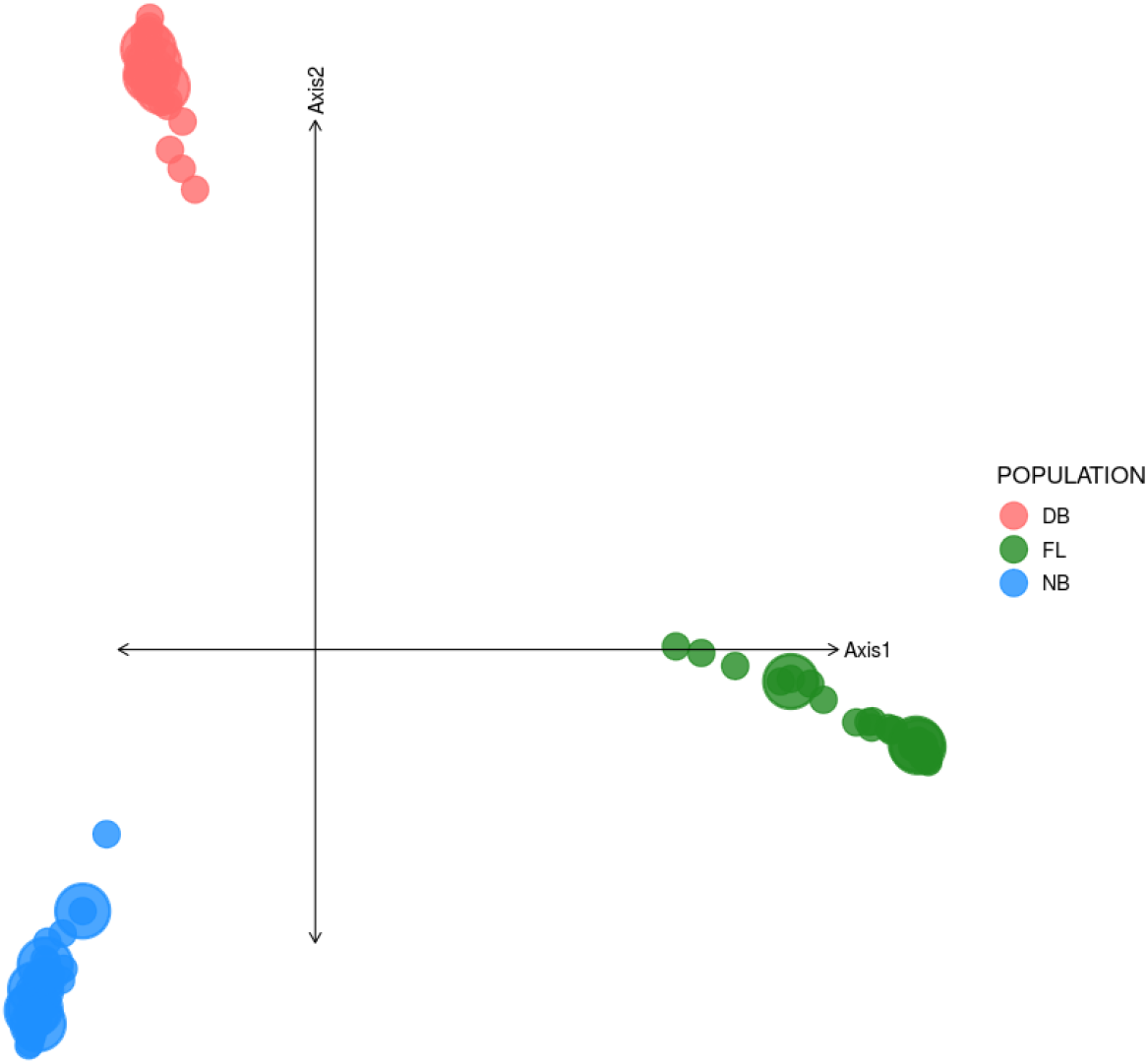
Principal component analysis on 100 individuals based on 2,180 diagnostic SNPs. Individuals are colored by lineage (DB: Delta Largemouth Bass, FL: Florida Bass, NB: northern Largemouth Bass). The top 500 SNPs based on PC loadings were used to design the 73 SNP diagnostic panel. PC1: 72.7% of variation, PC2: 13.3% of variation.

**Supp. Figure 3.**
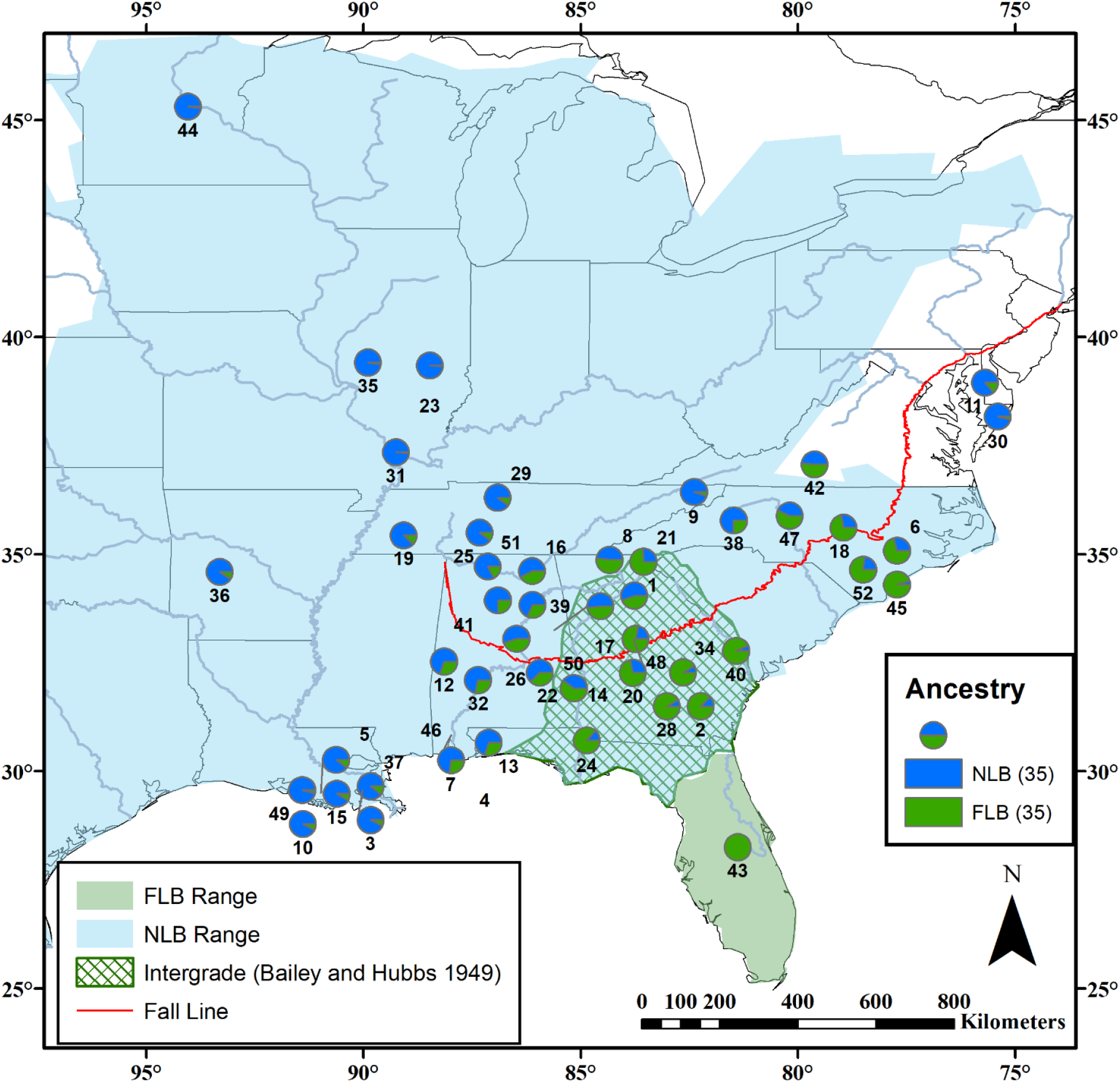
Sampling sites of largemouth bass assayed with 35 SNP panel diagnostic for northern Largemouth Bass (NLB) and Florida Bass (FLB). Populations are labelled as in Supp. File 2 and represented by pie graphs showing the mean estimated ancestry proportions for the two bass lineages based on counting the number of FLB and NLB alleles. Accepted native ranges of NLB and FLB are shown, as well as the NLB–FLB hybrid zone as described by Bailey & Hubbs, (1949). Map data from ESRI; FLB and NLB native ranges from Taylor et al., (2019).

**Supp. File 2.** Sampling sites assayed with 73 SNP panel and 35 SNP FLB–NLB panel. Numbers correspond to labels in Figure 1 and Supp. Figure 3, ancestry proportions are based on STRUCTURE results (K=3), using reference individuals.

**Supp. File 3.** Primer sequences (forward, reverse, and extension) designed to genotype two panels (52 and 39 SNPs using iPLEX PCR on the Agena MassARRAY System. 18 SNPs failed QC and are noted, however all primers must be included when genotyping with the MassARRAY platform and filtered after.

## References

Alford, J., & Jackson, D. (2009). Effects of Stocking Adult Largemouth Bass to Enhance Fisheries Recovery in Pascagoula River Floodplain Lakes Impacted by Hurricane Katrina. Proc. Annu. Conf. SEAFWA, 104–110.

Armstrong, D. L., Tucker, W. H., & Spray, R. G. (2000). Mobile delta management report 1999. Alabama Game and Fish.

Avise, J. C., & Saunders, N. C. (1984). Hybridization and introgression among species of sunfish (Lepomis): analysis by mitochondrial DNA and allozyme markers. Genetics, 108(1), 237–255.

Bagley, J. C., Mayden, R. L., Roe, K. J., Holznagel, W., & Harris, P. M. (2011). Congeneric phylogeographical sampling reveals polyphyly and novel biodiversity within black basses (Centrarchidae: Micropterus). Biological Journal of the Linnean Society, 104(2), 346–363.

Bailey, R. M., & Hubbs, C. L. (1949). The black basses (Micropterus) of Florida with description of a new species. University of Michigan Museum of Zoology.

Baker, W. H., Blanton, R. E., & Johnston, C. E. (2013). Diversity within the Redeye Bass, Micropterus coosae (Perciformes: Centrarchidae) species group, with descriptions of four new species. Zootaxa, 3635, 379–401.

Bangs, M. R., Oswald, K. J., Greig, T. W., Leitner, J. K., Rankin, D. M., & Quattro, J. M. (2018). Introgressive hybridization and species turnover in reservoirs: A case study involving endemic and invasive basses (Centrarchidae: Micropterus) in southeastern North America. Conservation Genetics, 19(1), 57–69.

Barthel, B. L., Lutz-Carrillo, D. J., Norberg, K. E., Porak, W. F., Tringali, M. D., Kassler, T. W., Johnson, W. E., Readel, A. M., Krause, R. A., & Philipp, D. P. (2010). Genetic relationships among populations of Florida Bass. Transactions of the American Fisheries Society, 139(6), 1615–1641.

Barton, N. H., & Hewitt, G. M. (1985). Analysis of hybrid zones. Annual Review of Ecology and Systematics, 16(1), 113–148.

Bermingham, E., & Avise, J. C. (1986). Molecular zoogeography of freshwater fishes in the southeastern United States. Genetics, 113(4), 939–965.

Bolnick, D. I. (2009). Hybridization and speciation in centrarchids. In S. J. Cooke & D. P. Philipp (Eds.), Centrarchid Fishes: Diversity, Biology and Conservation (pp. 39–69). Wiley-Blackwell Scientific Publications.

Bunch, A. J., Greenlee, R. S., & Odenkirk, J. (2017). Evaluation of Largemouth Bass Supplemental Stocking on a Virginia Coastal River.

Chhatre, V. E., Evans, L. M., DiFazio, S. P., & Keller, S. R. (2018). Adaptive introgression and maintenance of a trispecies hybrid complex in range-edge populations of Populus. Molecular Ecology, 27(23), 4820–4838.

Cooke, S. J., Kassler, T. W., & Philipp, D. P. (2001). Physiological performance of largemouth bass related to local adaptation and interstock hybridization: implications for conservation and management. Journal of Fish Biology, 59(sa), 248–268.

Cooke, S. J., & Philipp, D. P. (2006). Hybridization among divergent stocks of largemouth bass (Micropterus salmoides) results in altered cardiovascular performance: the influence of genetic and geographic distance. Physiological and Biochemical Zoology: PBZ, 79(2), 400–410.

Dakin, E. E., Porter, B. A., Freeman, B. J., & Long, J. M. (2015). Hybridization threatens shoal bass populations in the Upper Chattahoochee River Basin: Chapter 37. American Fisheries Society Southern Division Symposium 82, 491–502.

Danecek, P., Auton, A., Abecasis, G., Albers, C. A., Banks, E., DePristo, M. A., Handsaker, R. E., Lunter, G., Marth, G. T., Sherry, S. T., McVean, G., Durbin, R., & 1000 Genomes Project Analysis Group. (2011). The variant call format and VCFtools. Bioinformatics, 27(15), 2156–2158.

DeVries, D. R., Wright, R. A., Glover, D. C., Farmer, T. M., Lowe, M. R., Norris, A. J., & Peer, A. C. (2015). Largemouth Bass in coastal estuaries: a comprehensive study from the Mobile-Tensaw River Delta, Alabama. Black Bass Diversity, Multidisciplinary Science for Conservation, 82, 297–310.

Dyke, A. S., & Prest, V. K. (1987). Late Wisconsinan and Holocene history of the Laurentide ice sheet. Géographie Physique et Quaternaire, 41(2), 237.

Earl, D. A., & vonHoldt, B. M. (2012). STRUCTURE HARVESTER: a website and program for visualizing STRUCTURE output and implementing the Evanno method. Conservation Genetics Resources, 4(2), 359–361.

Edgar, R. C. (2004). MUSCLE: multiple sequence alignment with high accuracy and high throughput. Nucleic Acids Research, 32(5), 1792–1797.

Fitzpatrick, B. M. (2012). Estimating ancestry and heterozygosity of hybrids using molecular markers. BMC Evolutionary Biology, 12, 131.

Fitzpatrick, B. M., Johnson, J. R., Kump, D. K., Shaffer, H. B., Smith, J. J., & Voss, S. R. (2009). Rapid fixation of non-native alleles revealed by genome-wide SNP analysis of hybrid tiger salamanders. BMC Evolutionary Biology, 9, 176.

Fitzpatrick, B. M., Rya N, M. E., Johnson, J. R., Corush, J., & Carter, E. T. (2015). Hybridization and the species problem in conservation. Current Zoology, 61(1), 206–216.

Gabriel, S., Ziaugra, L., & Tabbaa, D. (2009). SNP genotyping using the Sequenom MassARRAY iPLEX platform. Current Protocols in Human Genetics, 60, 2.12.1–2.12.18.

Glover, D. C., DeVries, D. R., & Wright, R. A. (2012). Effects of temperature, salinity and body size on routine metabolism of coastal Largemouth Bass Micropterus salmoides. Journal of Fish Biology, 81(5), 1463–1478.

Glover, D. C., DeVries, D. R., & Wright, R. A. (2013). Growth of largemouth bass in a dynamic estuarine environment: an evaluation of the relative effects of salinity, diet, and temperature. Canadian Journal of Fisheries and Aquatic Sciences. Journal Canadien Des Sciences Halieutiques et Aquatiques, 70(3), 485–501.

Gompert, Z., Mandeville, E. G., & Buerkle, C. A. (2017). Analysis of population genomic data from hybrid zones. Annual Review of Ecology, Evolution, and Systematics, 48(1), 207–229.

Gompert, Z., Parchman, T. L., & Buerkle, C. A. (2012). Genomics of isolation in hybrids. Philosophical Transactions of the Royal Society of London. Series B, Biological Sciences, 367(1587), 439–450.

Goudet, J., & Jombart, T. (2015). hierfstat: Estimation and Tests of Hierarchical F-Statistics (Version 0.04-22). https://CRAN.R-project.org/package=hierfstat

Guindon, S., Dufayard, J.-F., Lefort, V., Anisimova, M., Hordijk, W., & Gascuel, O. (2010). New algorithms and methods to estimate maximum-likelihood phylogenies: assessing the performance of PhyML 3.0. Systematic Biology, 59(3), 307–321.

Hallerman, E. M., Smitherman, R. O., Reed, R. B., Tucker, W. H., & Dunham, R. A. (1986). Biochemical genetics of Largemouth bass in mesosaline and freshwater areas of the Alabama river system. Transactions of the American Fisheries Society, 115(1), 15–20.

Hargrove, J. S., Rogers, M. W., Kacmar, P. T., & Black, P. (2019). A statewide evaluation of Florida bass genetic introgression in Tennessee. North American Journal of Fisheries Management, 39(4), 637–651.

Harrison, R. G., & Larson, E. L. (2014). Hybridization, introgression, and the nature of species boundaries. The Journal of Heredity, 105 Suppl 1, 795–809.

Hijmans, R. J., Williams, E., & Vennes, C. (2016). Geosphere: spherical trigonometry. R package.

Hoang, D. T., Chernomor, O., von Haeseler, A., Minh, B. Q., & Vinh, L. S. (2018). UFBoot2: Improving the Ultrafast Bootstrap Approximation. Molecular Biology and Evolution, 35(2), 518–522.

Jombart, T., & Ahmed, I. (2011). adegenet 1.3-1: new tools for the analysis of genome-wide SNP data. Bioinformatics, 27(21), 3070–3071.

Kalyaanamoorthy, S., Minh, B. Q., Wong, T. K. F., von Haeseler, A., & Jermiin, L. S. (2017). ModelFinder: fast model selection for accurate phylogenetic estimates. Nature Methods, 14(6), 587–589.

Kassler, T., Koppelman, J., Near, T., Dillman, C., Levengood, J., Swofford, D., VanOrman, J., Claussen, J., & Phillip, D. (2002). Molecular and morphological analyses of the black basses: implications for taxonomy and conservation. American Fisheries Society Symposium, 31, 291–322.

Keck, B. P., & Near, T. J. (2009). Geographic and temporal aspects of mitochondrial replacement in Nothonotus darters (Teleostei: Percidae: Etheostomatinae). Evolution; International Journal of Organic Evolution, 64(5), 1410–1428.

Kocher, T. D., Conroy, J. A., McKaye, K. R., Stauffer, J. R., & Lockwood, S. F. (1995). Evolution of NADH dehydrogenase subunit 2 in east African cichlid fish. Molecular Phylogenetics and Evolution, 4(4), 420–432.

Kopelman, N. M., Mayzel, J., Jakobsson, M., Rosenberg, N. A., & Mayrose, I. (2015). CLUMPAK: A program for identifying clustering modes and packaging population structure inferences across K. Molecular Ecology Resources, 15(5), 1179–1191.

Kumar, S., Stecher, G., Li, M., Knyaz, C., & Tamura, K. (2018). MEGA X: Molecular Evolutionary Genetics Analysis across Computing Platforms. Molecular Biology and Evolution, 35(6), 1547–1549.

Largiadèr, C. R. (2007). Hybridization and Introgression Between Native and Alien Species. In W. Nentwig (Ed.), Biological Invasions (pp. 275–292). Springer Berlin Heidelberg.

Li, C., Gowan, S., Anil, A., Beck, B. H., Thongda, W., Kucuktas, H., Kaltenboeck, L., & Peatman, E. (2015). Discovery and validation of gene-linked diagnostic SNP markers for assessing hybridization between Largemouth bass (Micropterus salmoides) and Florida bass (M. floridanus). Molecular Ecology Resources, 15(2), 395–404.

Li, H., & Durbin, R. (2009). Fast and accurate short read alignment with Burrows–Wheeler transform. Bioinformatics.

Li, H., Handsaker, B., Wysoker, A., Fennell, T., Ruan, J., Homer, N., Marth, G., Abecasis, G., Durbin, R., & 1000 Genome Project Data Processing Subgroup. (2009). The Sequence Alignment/Map format and SAMtools. Bioinformatics, 25(16), 2078–2079.

Lutz-Carrillo, D. J., Nice, C. C., Bonner, T. H., Forstner, M. R. J., & Fries, L. T. (2006). Admixture analysis of Florida largemouth bass and Northern largemouth bass using microsatellite loci. Transactions of the American Fisheries Society, 135(3), 779–791.

Maceina, M. J., & Murphy, B. R. (1992). Stocking Florida Largemouth Bass outside its native range. Transactions of the American Fisheries Society, 121(5), 686–691.

Maceina, M. J., Murphy, B. R., & Isely, J. J. (1988). Factors regulating Florida Largemouth bass stocking success and hybridization with northern Largemouth bass in Aquilla lake, Texas. Transactions of the American Fisheries Society, 117(3), 221–231.

Malde, K., Seliussen, B. B., Quintela, M., Dahle, G., Besnier, F., Skaug, H. J., Øien, N., Solvang, H. K., Haug, T., Skern-Mauritzen, R., Kanda, N., Pastene, L. A., Jonassen, I., & Glover, K. A. (2017). Whole genome resequencing reveals diagnostic markers for investigating global migration and hybridization between minke whale species. BMC Genomics, 18(1), 76.

McDonald, D. B., Parchman, T. L., Bower, M. R., Hubert, W. A., & Rahel, F. J. (2008). An introduced and a native vertebrate hybridize to form a genetic bridge to a second native species. Proceedings of the National Academy of Sciences of the United States of America, 105(31), 10837–10842.

Mendes, S. L., Machado, M. P., Coelho, M. M., & Sousa, V. C. (2021). Genomic data and multispecies demographic modelling uncover past hybridization between currently allopatric freshwater species. In bioRxiv (p. 585687). https://doi.org/10.1101/585687

Meyer, K. A., Kennedy, P., High, B., & Campbell, M. R. (2017). Distinguishing Yellowstone Cutthroat Trout, Rainbow Trout, and Hybrids by Use of Field-Based Phenotypic Characteristics. North American Journal of Fisheries Management, 37(2), 456–466.

Near, T. J., Kassler, T. W., Koppelman, J. B., Dillman, C. B., & Philipp, D. P. (2003). Speciation in north American black basses, Micropterus (Actinopterygii: Centrarchidae). Evolution, 57(7), 1610.

Near, T. J., & Kim, D. (2021). Phylogeny and time scale of diversification in the fossil-rich sunfishes and black basses (Teleostei: Percomorpha: Centrarchidae). Molecular Phylogenetics and Evolution, 161, 107156.

Nei, M., & Chesser, R. K. (1983). Estimation of fixation indices and gene diversities. Annals of Human Genetics, 47(3), 253–259.

Nei, M., Maruyama, T., & Chakraborty, R. (1975). The bottleneck effect and genetic variability in populations. Evolution, 29(1), 1–10.

Nevado, B., Fazalova, V., Backeljau, T., Hanssens, M., & Verheyen, E. (2011). Repeated unidirectional introgression of nuclear and mitochondrial DNA between four congeneric Tanganyikan cichlids. Molecular Biology and Evolution, 28(8), 2253–2267.

Nguyen, L.-T., Schmidt, H. A., von Haeseler, A., & Minh, B. Q. (2015). IQ-TREE: a fast and effective stochastic algorithm for estimating maximum-likelihood phylogenies. Molecular Biology and Evolution, 32(1), 268–274.

Novembre, J., Williams, R., Pourreza, H., Wang, Y., & Carbonetto, P. (2018). PCAviz: Visualizing Principal Components Analysis (Version 0.3-29). http://github.com/NovembreLab/PCAviz

Ortiz, E. M. (2019). vcf2phylip v2.0: convert a VCF matrix into several matrix formats for phylogenetic analysis. https://doi.org/10.5281/zenodo.2540861

Paradis, E. (2010). pegas: an R package for population genetics with an integrated–modular approach. Bioinformatics, 26(3), 419–420.

Peñaloza-Ramírez, J. M., González-Rodríguez, A., Mendoza-Cuenca, L., Caron, H., Kremer, A., & Oyama, K. (2010). Interspecific gene flow in a multispecies oak hybrid zone in the Sierra Tarahumara of Mexico. Annals of Botany, 105(3), 389–399.

Philipp, D. P., Childers, W. F., & Whitt, G. S. (1983). A biochemical genetic evaluation of the northern and Florida subspecies of Largemouth bass. Transactions of the American Fisheries Society, 112(1), 1–20.

Philipp, D. P., & Claussen, J. E. (1995). Fitness and performance differences between two stocks of largemouth bass from different river drainages within Illinois. American Fisheries Society Symposium, 15, 236–243.

Pickrell, J. K., & Pritchard, J. K. (2012). Inference of population splits and mixtures from genome-wide allele frequency data. PLoS Genetics, 8(11), e1002967.

Piganeau, G., Gardner, M., & Eyre-Walker, A. (2004). A broad survey of recombination in animal mitochondria. Molecular Biology and Evolution, 21(12), 2319–2325.

Puritz, J. B., Hollenbeck, C. M., & Gold, J. R. (2014). dDocent: A RADseq, variant-calling pipeline designed for population genomics of non-model organisms. PeerJ, 2, e431.

Rainer, D. (2010, July). Delta bass: fisheries biologists are developing a new strain of bass brood stock to release in the Mobile-Tensaw delta. Outdoor Alabama, 12–14.

Rhymer, J. M., & Simberloff, D. (1996). Extinction by hybridization and introgression. Annual Review of Ecology and Systematics, 27(1), 83–109.

Rochette, N. C., Rivera-Colón, A. G., & Catchen, J. M. (2019). Stacks 2: Analytical methods for paired-end sequencing improve RADseq-based population genomics. Molecular Ecology, 28(21), 4737–4754.

Seyoum, S., Barthel, B. L., Tringali, M. D., Davis, M. C., Schmitt, S. L., Bellotti, P S., & Porak, W. F. (2013). Isolation and characterization of eighteen microsatellite loci for the largemouth bass, Micropterus salmoides, and cross amplification in congeneric species. Conservation Genetics Resources, 5(3), 697–701.

Simone, D., Calabrese, F. M., Lang, M., Gasparre, G., & Attimonelli, M. (2011). The reference human nuclear mitochondrial sequences compilation validated and implemented on the UCSC genome browser. BMC Genomics, 12, 517.

Stepien, C. A., Sepulveda-Villet, O. J., & Haponski, A. E. (2015). Comparative Genetic Diversity, Population Structure, and Adaptations of Walleye and Yellow Perch Across North America. In P. Kestemont, K. Dabrowski, & R. C. Summerfelt (Eds.), Biology and Culture of Percid Fishes: Principles and Practices (pp. 643–689). Springer Netherlands.

Swift, C. C., Gilbert, C. R., Bortone, S. A., Burgess, G. H., & Yerger, R. W. (1986). Zoogeography of the freshwater fishes of the southeastern United States: Savannah River to Lake Pontchartrain. The Zoogeography of North American Freshwater Fishes.

Swingle, H. A., & Bland, D. G. (1974). A Study of the Fishes of the Coastal Water-courses of Alabama. Alabama Marine Resources Laboratory.

Swingle, W. E., Spencer, S. L., & Scott, T. M., Jr. (1966). Statistics on the sport fishery of the Mobile Delta during the period of July 1, 1963 to June 30, 1964. Proceedings of the Southeastern Association of Game and Fish Commissioners, 19, 439–446.

Taylor, A. T., Long, J. M., Tringali, M. D., & Barthel, B. L. (2019). Conservation of black bass diversity: An emerging management paradigm. Fisheries, 44(1), 20–36.

Taylor, A. T., Tringali, M. D., O’Rouke, P M., & Long, J. M. (2018). Shoal bass hybridization in the Chattahoochee River Basin near Atlanta, Georgia. Journal of Southeastern Association of Fish and Wildlife Agencies, 5, 1–9.

Taylor, S. A., Larson, E. L., & Harrison, R. G. (2015). Hybrid zones: windows on climate change. Trends in Ecology & Evolution, 30(7), 398–406.

Thongda, W., Lewis, M., Zhao, H., Bowen, B., Lutz-Carrillo, D. J., Peoples, B. K., & Peatman, E. (2020). Species-diagnostic SNP markers for the black basses (Micropterus spp.): a new tool for black bass conservation and management. Conservation Genetics Resources, 12, 319–328.

Travnichek, V. H., Maceina, M. J., Smith, S. M., & Dunham, R. A. (1996). Natural hybridization between Black and White Crappies (Pomoxis) in 10 Alabama Reservoirs. The American Midland Naturalist, 135(2), 310–316.

Truett, G. E., Heeger, P., Mynatt, R. L., Truett, A. A., Walker, J. A., & Warman, M. L. (2000). Preparation of PCR-quality mouse genomic DNA with hot sodium hydroxide and tris (HotSHOT). BioTechniques, 29(1), 52, 54.

Tucker, W. H. (1985). Age and growth of Largemouth Bass in the Mobile delta. The Journal of the Alabama Academy of Science, 56(2), 65–70.

Vähä, J.-P., & Primmer, C. R. (2006). Efficiency of model-based Bayesian methods for detecting hybrid individuals under different hybridization scenarios and with different numbers of loci. Molecular Ecology, 15(1), 63–72.

Viard, F., Riginos, C., & Bierne, N. (2020). Anthropogenic hybridization at sea: three evolutionary questions relevant to invasive species management. Philosophical Transactions of the Royal Society of London. Series B, Biological Sciences, 375(1806), 20190547.

Warren, M. L., Angermeier, P L., Burr, B. M., & Haag, W. R. (1997). Decline of a diverse fish fauna: patterns of imperilment and protection in the southeastern United States. In G. W. Benz & D. E. Collins (Eds.), Aquatic Fauna in Peril: The Southeastern Perspective. Southeast Aquatic Research Institute.

Weir, B. S., & Cockerham, C. C. (1984). Estimating F-statistics for the analysis of population structure. Evolution, 38(6), 1358–1370.

Wilk, R. J., & Horth, L. (2016). A genetically distinct hybrid zone occurs for two globally invasive mosquito fish species with striking phenotypic resemblance. Ecology and Evolution, 6(23), 8375–8388.

Woodruff, D. S. (1973). Natural Hybridization and Hybrid Zones. Systematic Biology, 22(3), 213–218.

Young, J. L., Bornik, Z. B., Marcotte, M. L., Charlie, K. N., Wagner, G. N., Hinch, S. G., & Cooke, S. J. (2006). Integrating physiology and life history to improve fisheries management and conservation. Fish and Fisheries, 7(4), 262–283.

Yu, G., Smith, D. K., Zhu, H., Guan, Y., & Lam, T. T. (2017). Ggtree: An r package for visualization and annotation of phylogenetic trees with their covariates and other associated data. Methods in Ecology and Evolution / British Ecological Society, 8(1), 28–36.

Zhao, H., Li, C., Hargrove, J. S., Bowen, B. R., Thongda, W., Zhang, D., Mohammed, H., Beck, B. H., Austin, J. D., & Peatman, E. (2018). SNP marker panels for parentage assignment and traceability in the Florida bass (Micropterus floridanus). Aquaculture, 485, 30–38.

Zhao, H., Silliman, K., Lewis, M., Johnson, S., Kratina, G., Rider, S. J., Stepien, C. A., Hallerman, E. M., Beck, B., Fuller, A., & Peatman, E. (2020). SNP analyses highlight a unique, imperiled southern walleye (Sander vitreus) in the Mobile River Basin. Canadian Journal of Fisheries and Aquatic Sciences, 77, 1366–1378.

